# Coupled single-cell CRISPR screening and epigenomic profiling reveals causal gene regulatory networks

**DOI:** 10.1101/414870

**Authors:** Adam J. Rubin, Kevin R. Parker, Ansuman T. Satpathy, Yanyan Qi, Beijing Wu, Alvin J. Ong, Maxwell R. Mumbach, Andrew L. Ji, Daniel S. Kim, Seung Woo Cho, Brian J. Zarnegar, William J. Greenleaf, Howard Y. Chang, Paul A. Khavari

## Abstract

Here we present Perturb-ATAC, a method which combines multiplexed CRISPR interference or knockout with genome-wide chromatin accessibility profiling in single cells, based on the simultaneous detection of CRISPR guide RNAs and open chromatin sites by assay of transposase-accessible chromatin with sequencing (ATAC-seq). We applied Perturb-ATAC to transcription factors (TFs), chromatin-modifying factors, and noncoding RNAs (ncRNAs) in ∼4,300 single cells, encompassing more than 63 unique genotype-phenotype relationships. Perturb-ATAC in human B lymphocytes uncovered regulators of chromatin accessibility, TF occupancy, and nucleosome positioning, and identified a hierarchical organization of TFs that govern B cell state, variation, and disease-associated *cis*-regulatory elements. Perturb-ATAC in primary human epidermal cells revealed three sequential modules of *cis*-elements that specify keratinocyte fate, orchestrated by the TFs JUNB, KLF4, ZNF750, CEBPA, and EHF. Combinatorial deletion of all pairs of these TFs uncovered their epistatic relationships and highlighted genomic co-localization as a basis for synergistic interactions. Thus, Perturb-ATAC is a powerful and general strategy to dissect gene regulatory networks in development and disease.

**Highlights:**

1. A new method for simultaneous measurement of CRISPR perturbations and chromatin state in single cells.
2. Perturb-ATAC reveals regulatory factors that control *cis*-element accessibility, *trans*-factor occupancy, and nucleosome positioning.
3. Perturb-ATAC reveals regulatory modules of coordinated *trans*-factor activity in B lymphoblasts.
4. Keratinocyte differentiation is orchestrated by synergistic activities of co-binding TFs on *cis*-elements.

## Introduction

Gene expression in eukaryotic organisms is regulated by a complex interplay of thousands of *trans*-acting regulatory factors and millions of *cis*-acting DNA elements (Roadmap Epigenomics Consortium et al., 2015). These precise epigenetic interactions are required to establish gene expression patterns that underlie the development of distinct cell types, the interactions of these cell types in multicellular tissues, and their responses to a wide array of environmental stimuli (Rada-Iglesias et al., 2011; Kaikkonen et al., 2013). However, it has remained a significant challenge to dissect the molecular contributions of each *trans*-factor or *cis*-element to establishing the global chromatin state, since epigenetic assays: 1) require large amounts of starting cellular material, 2) are time-consuming to perform, and 3) are difficult to couple with genetic perturbations at scale. Therefore, prior studies have largely focused on measuring the effects of single genetic perturbations on the chromatin state in bulk cell populations or large tissue samples.

We recently developed the assay for transposase-accessible chromatin with sequencing (ATAC-seq), which utilizes a hyperactive transposase (Tn5) to measure regulatory DNA elements by direct transposition of sequencing adaptors into regions of accessible chromatin in a cell (Buenrostro et al., 2013). The resulting ATAC-seq profile can provide insight into several layers of epigenetic regulation, including the identification of enhancer and promoter sequences genome-wide with basepair (bp)-resolution, the precise positioning of nucleosomes, and the inference of transcription factors bound to each site through DNA foot-printing of transposase-inaccessible regions (Buenrostro et al., 2013; Schep et al., 2015, 2017). The accumulation of these layers of epigenetic information, all read out as chromatin accessibility signal from a single assay, provides a rich view of the chromatin state, which can then be interrogated in the setting of genetic perturbations. Importantly, ATAC-seq can be performed in single cells (scATAC-seq), which can be utilized in several ways (Buenrostro et al., 2015; Cusanovich et al., 2015). First, scATAC-seq profiles can identify cell-to-cell variability and intermediate epigenomic phenotypes that are obscured by bulk measurements (Cusanovich et al., 2018; Satpathy et al., 2018; Buenrostro et al., 2018). Second, scATAC-seq profiles can be generated in high-throughput, from 100s to 1000s of single cells in one experiment. Finally, scATAC-seq profiles can be paired with orthogonal measurements of RNA or protein expression in the same cell (Satpathy et al., 2018; Chen et al., 2018a; Liu et al., 2018).

Similar advances in the ability to measure transcriptomes in single cells have recently enabled high-throughput genetic screens coupled with simultaneous phenotypic measurements. Standard genetic screens are typically performed by generating a diverse set of mutations in a population of cells and then selecting competitive outgrowth of single cellular phenotypes, such as survival or the expression of specific surface markers. Following selection, mutations that are enriched or depleted in the population are identified and further individually characterized to determine genome-wide effects. However, this screening strategy is challenging to perform for many complex cellular phenotypes that may not perfectly track with a known selection protocol. Single-cell transcriptome profiling enabled an alternative strategy, whereby the effects of perturbations on phenotypes are scored in high-throughput using paired sgRNA and phenotype assessment in single cells (Dixit et al., 2016; Adamson et al., 2016; Jaitin et al., 2016; Datlinger et al., 2017). These studies have used single-cell transcriptome profiling to identify gene targets, gene signatures, and cell states that are impacted by each perturbation in a given cell. In this study, we further develop this concept of genetic screening by measuring the effects of perturbations on chromatin accessibility in single cells genome-wide (**Figure 1**). This method, termed perturbation-indexed single-cell ATAC-seq (Perturb-ATAC) enables deep profiling of the regulatory landscape, including identification of chromatin accessibility at enhancers and promoters, TF binding to DNA, and nucleosome positioning – genomic features that can only be measured using epigenetic profiling.

**Figure 1.**
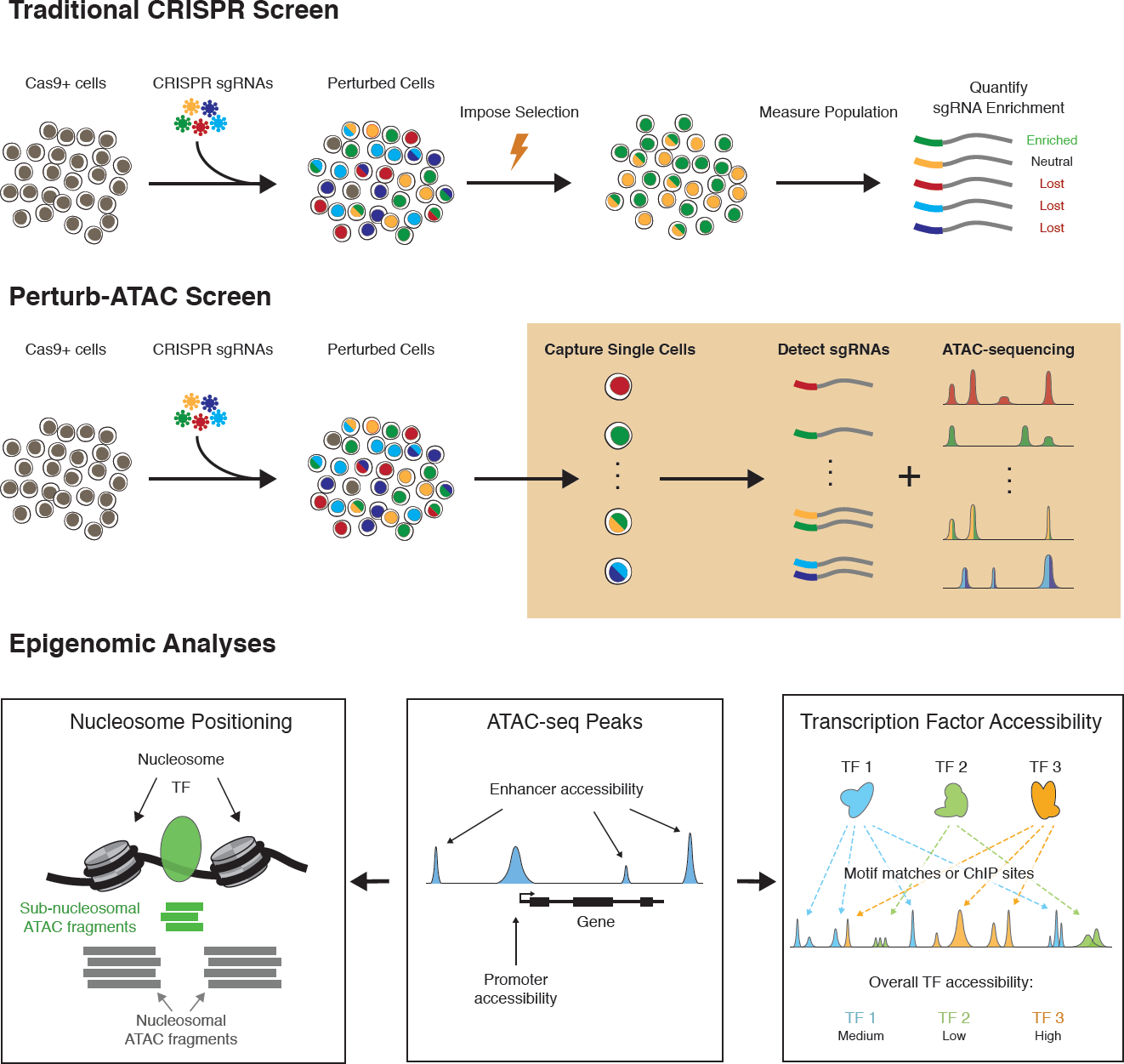
Perturb-ATAC: simultaneous CRISPR guide RNA detection and epigenome profiling for high complexity genetic screens. Top: Schematic describing traditional CRISPR screens. Middle: Perturb-ATAC workflow. Bottom: Overview of classes of biological questions that can be interrogated from Perturb-ATAC data.

Perturb-ATAC is an experimental and computational methodology that enables high-throughput profiling of chromatin accessibility combined with genetic perturbations in single cells, by simultaneously measuring Clustered Regularly Interspaced Short Palindromic Repeats (CRISPR) single guide RNA (sgRNA) sequences and scATAC-seq profiles in each cell. To demonstrate the performance and utility of Perturb-ATAC, we performed this method in 2,936 immortalized B lymphoblasts and 1,356 primary human keratinocytes, encompassing a total of 63 genotype-phenotype relationships. Analysis of a CRISPR-interference (CRISPRi) screen in GM12878 lymphoblasts identified *trans*-factors that controlled several layers of epigenetic regulation associated with the B cell state. scATAC-seq in primary human epidermal cells identified three regulatory modules that accompanied keratinocyte differentiation, and Perturb-ATAC using a CRISPR-deletion screen revealed key regulators of each module, including the transcription factors JUNB, KLF4, ZNF750, CEBPA, and EHF. We further mapped epistatic regulatory relationships between these TFs using multiplexed perturbations in single cells. Dual perturbations uncovered a surprising degree of synergy between factors, and further analysis of these relationships showed that genomic co-localization and co-expression of TFs may be predictors of genetic interaction. Altogether, these results establish Perturb-ATAC as a new tool for studying relationships between the factors that control chromatin states using high-throughput, high-complexity single-cell screens.

## Results

### Perturb-ATAC: simultaneous CRISPR guide detection and epigenome profiling in single cells

We implemented Perturb-ATAC by adapting two prior technologies to measure single-cell chromatin accessibility using ATAC-seq and to simultaneously measure targeted RNA sequences and ATAC-seq in single cells using a microfluidic platform (Buenrostro et al., 2015; Satpathy et al., 2018). In the Perturb-ATAC protocol, single cells are first individually captured on the Integrated Fluidics Circuit (IFC; Fluidigm) in single-cell chambers and then subjected to cell lysis and DNA transposition with the hyperactive Tn5 enzyme (**Figure 2A**). After transposition, Tn5 is released from open chromatin fragments, and CRISPR sgRNAs or sgRNA-identifying barcodes within each chamber are subjected to reverse transcription (RT) using primers targeting regions flanking each sequence. Immediately after RT, 5’ ends of ATAC-seq fragments are extended and all chamber contents are amplified by PCR. Single-cell libraries are then collected and sgRNA or ATAC amplicons are further amplified separately with cell-identifying barcoded primers, pooled, and sequenced on a high-throughput sequencing instrument.

**Figure 2.**
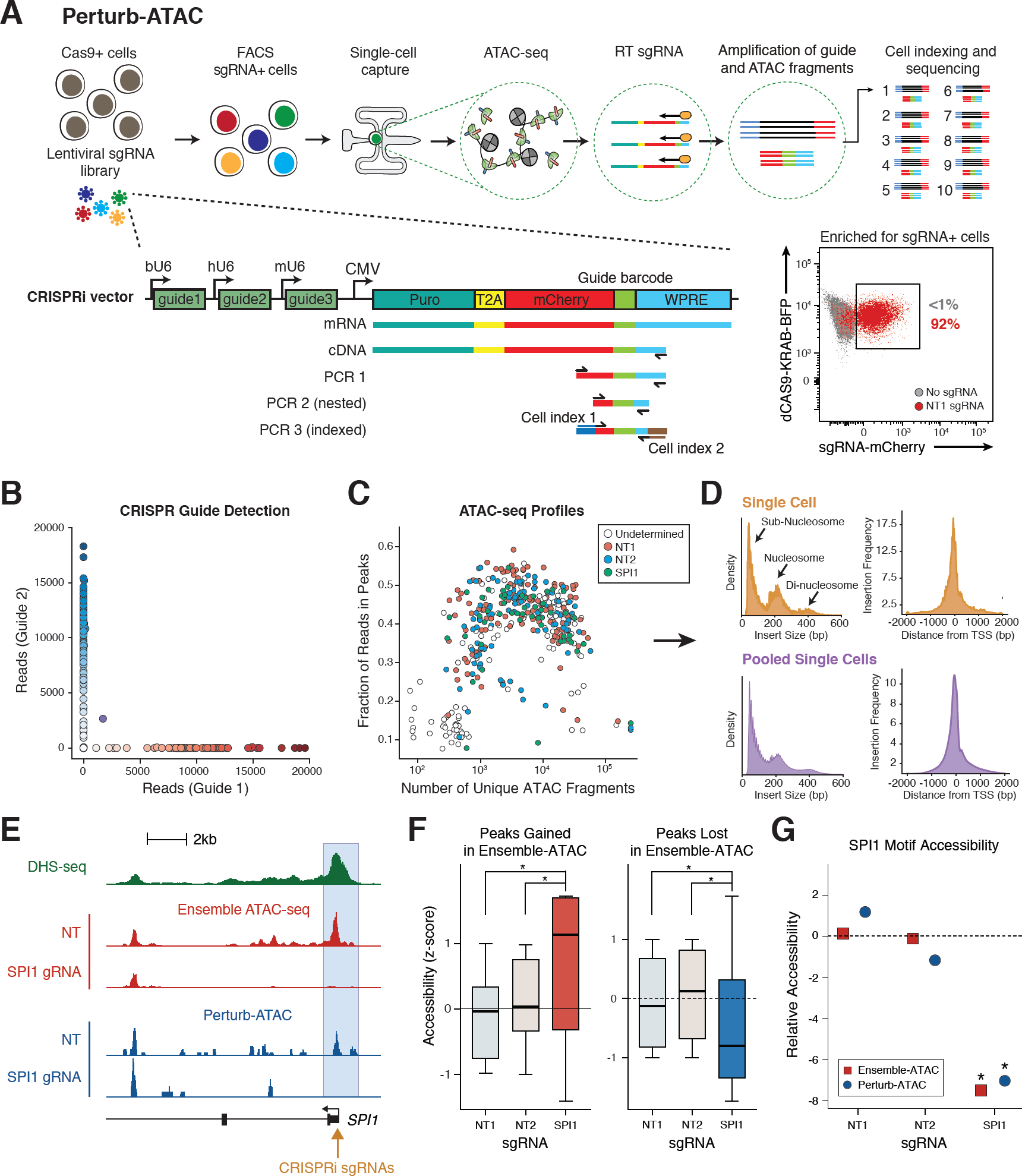
Perturb-ATAC identifies guide RNA barcodes and expected chromatin phenotypes in single cells. **(a)** Schematic of Perturb-ATAC protocol, lentiviral construct, and generation of sequencing library for guide RNA detection. **(b)** Scatter plot of guide barcode reads from pool of cells transduced with one of two guide constructs. **(c)** Scatter plot of ATAC fragments and the fraction of ATAC fragments in peak regions for each cell. Labels indicating guide barcode detection in each cell are shown. **(d)** Density histograms of ATAC fragment size distribution (left) indicating expected nucleosome phasing and relative frequency of ATAC insertions surrounding transcription start sites (right) in merged single cells (top) and bulk cells (bottom). **(e)** Genomic locus of *SPI1* gene, indicating DNase I hypersensitivity sequencing, bulk ATAC-seq, and Perturb-ATAC-seq. The SPI1 promoter region exhibits selective loss of accessibility in cells expressing SPI1 sgRNA. **(f)** Box plots of accessibility from merged single cells of individual genomic regions identified as altered in bulk ATAC-seq. * indicates p-value < 1e^-3^ by KS-test. **(g)** Relative accessibility of SPI1 motif-containing regions (z-score of relative activity of SPI1 motif versus all other genomic features). * indicates false discovery rate < 1e^-3^ by permutation test.

To initially test the performance of Perturb-ATAC, we adapted a CRISPR interference (CRISPRi) sgRNA vector to perform a simple mixing experiment with three populations of sgRNA-targeted cells (Adamson et al., 2016; Cho et al., 2018). We generated a sgRNA vector that included unique 22-basepair (bp) guide barcodes (GBCs) that corresponded to the identity of sgRNAs encoded by each vector (**Figures 2A and S1A**). Using this vector, we targeted immortalized B lymphoblasts stably expressing dCas9-KRAB with either non-human-genome-targeting (NT) sgRNAs (NT1 or NT2) or sgRNAs targeting the promoter of the transcription factor SPI1 (also known as PU.1), which is required for B cell development (Scott et al., 1994; McKercher et al., 1996; Scott et al., 1997). We then pooled cells post-transduction and performed Perturb-ATAC on 288 single cells to assess the fidelity of pairing GBC detection with the measurement of epigenetic phenotypes. To eliminate the likelihood of recombination of sgRNAs and GBCs between vectors in a pooled viral packaging experiment, as has been recently described (Adamson et al., 2018; Feldman et al., 2018; Hill et al., 2018; Xie et al., 2018), we performed all packaging steps in individual cultures and then pooled viruses for subsequent transduction. Following transduction, lentivirus-containing cells were enriched by FACS-purification of mCherry+ cells, and the identity of the sgRNAs delivered to each cell was read out through amplification of the GBC (**Figure 2A**).

We used stringent cutoffs to assign the presence or absence of a GBC in a given cell (**Figure S1B**). First, we counted the number of reads for each possible GBC in every cell and then adjusted these counts for sequencing depth to account for plate-to-plate variation during library preparation or sequencing (**Figures S1B and S1C**). Next, we set a minimum read cutoff of 1,000 GBC reads per cell to remove cells with low coverage, and removed cells with a high percentage of background reads (non-specific amplification reads; **Methods**). Finally, while this pilot experiment contained only single-sgRNA cells, in later experiments where cells could be either singly- or doubly-transduced, we set a cutoff based on the percent of GBC reads aligning to the second-most common GBC (single GBC < 1%; double GBC, > 5%; **Methods**). Importantly, the majority of cells passed each of these filtering steps, and analysis of cells passing filter in the mixing experiment showed that GBC reads in each cell were almost exclusively assigned to one GBC (NT1, NT2, or SPI1 sgRNAs), as expected; multiple GBCs were only detected in 9/309 cells. These results demonstrate that Perturb-ATAC consistently detects GBCs with high confidence and a low false positive rate in single cells (**Figures 2B, S1B and S1C**).

Next, we evaluated the quantity and quality of ATAC-seq reads in single cells assayed by Perturb-ATAC. High quality single-cell ATAC-seq profiles were obtained in 79.2% of cells in which GBCs were also detected (**Figure 2C**). Cells passing filter yielded an average of 11.33 × 10^3^ fragments mapping to the nuclear genome, and approximately 43.05% of reads were within peaks present in bulk GM12878 ATAC-seq profiles (**Figure 2C**). This fraction of reads in peaks is similar to previously published scATAC-seq as well as high-quality bulk ATAC-seq datasets (Buenrostro et al., 2015; Corces et al., 2017). Single-cell ATAC-seq reads recapitulated several characteristics of bulk ATAC-seq data, including insert-size periodicity from nucleosome protection of DNA and enrichment of ATAC-seq fragments at transcription start sites (TSS) (**Figure 2D**). These results indicate that GBC detection does not interfere with the generation of ATAC-seq libraries from single cells.

Finally, we asked whether single cells containing each GBC exhibited the expected ATAC-seq phenotype. ATAC-seq data can be analyzed at the level of individual regulatory DNA elements or at the level of TF activity genome-wide, computed from observed/expected ATAC-seq reads in TF binding sites in each cell. For individual regulatory elements, we first examined the promoter of the *SPI1* gene, where *SPI1*-targeted sgRNAs were expected to recruit dCas9-KRAB and alter the local chromatin state. In single cells receiving *SPI1* sgRNAs, we observed a selective loss of accessibility at the promoter, compared to cells receiving NT sgRNAs (**Figure 2E**). More broadly, *SPI1*-targeted single cells exhibited similar changes in accessibility (gain or loss) across all peaks that were changed in bulk ATAC-seq *SPI1*-targeting experiments (**Figure 2F**). For measurement of global TF activity, we determined the relative accessibility of all SPI1 motif-containing sites genome-wide (51,862 sites). We found that SPI1 sites globally exhibited a significant loss of accessibility in *SPI1*-targeted cells compared to NT cells (FDR < 1e-3, permutation test; **Methods**), and that the magnitude of loss in accessibility was comparable between single cells and independent bulk *SPI1*-targeting experiments (**Figure 2G**). Altogether, these findings demonstrate that Perturb-ATAC can simultaneously measure GBC sequences and ATAC-seq data in single cells with high fidelity.

### Perturb-ATAC identifies epigenomic functions of chromatin regulators, transcription factors, and noncoding RNAs in B cells

We next performed an expanded Perturb-ATAC screen to compare how broadly-expressed and lineage-specific *trans*-factors shape the chromatin landscape of B lymphoblasts. We targeted dCas9-KRAB-expressing GM12878 cells to generate 40 sgRNA genotypes in 2,627 single cells, derived from single- or combinatorial-targeting of the promoters of 12 *trans*-factors and 2 NT control sgRNAs. The 12 targeted *trans*-factors included transcription factors (EBF1, IRF8, NFKB1, RELA, and SPI1), chromatin modifiers (BRG1, DNMT3A, EZH2, and TET2), and noncoding RNAs (7SK, EBER1, and EBER2) that are expressed in B cells and have previously been shown to impact normal and neoplastic B cell development and function (Nutt and Kee, 2007; Lunning and Green, 2015). CRISPRi guide RNAs were designed to maximize knockdown efficiency and minimize off-target effects at mismatch loci (**Figure S2A-C**). To assess possible epistatic relationships between *trans*-factors, we transduced cells with single or multiple sgRNA constructs and distinguished cells receiving either one or two GBCs, as described above (**Figures 3A and 3B**). Similar to the mixing experiment, high quality single-cell ATAC-seq profiles were obtained in 85.3% of cells in which GBCs were also detected. Single cells passing filter yielded an average of 10.68 × 10^3^ fragments mapping to the nuclear genome, and approximately 43.85% of reads were within peaks present in bulk GM12878 ATAC-seq profiles (**Figure 3C**). The presence of ATAC reads mapping to any segment of the viral construct was extremely rare and did not appear to influence epigenomic profiles (**Methods**). Direct observation of accessibility at potential sgRNA mismatch loci indicated no evidence of off-target repression, highlighting that ATAC profiles reflect the effects of target gene repression (**Figure S2D**). Cells were processed across several microfluidic chips, and samples from each chip exhibited no noticeable bias in ATAC-seq signal (**Figure S2E**), consistent with production of technically sound datasets in these experiments for subsequent analysis.

**Figure 3.**
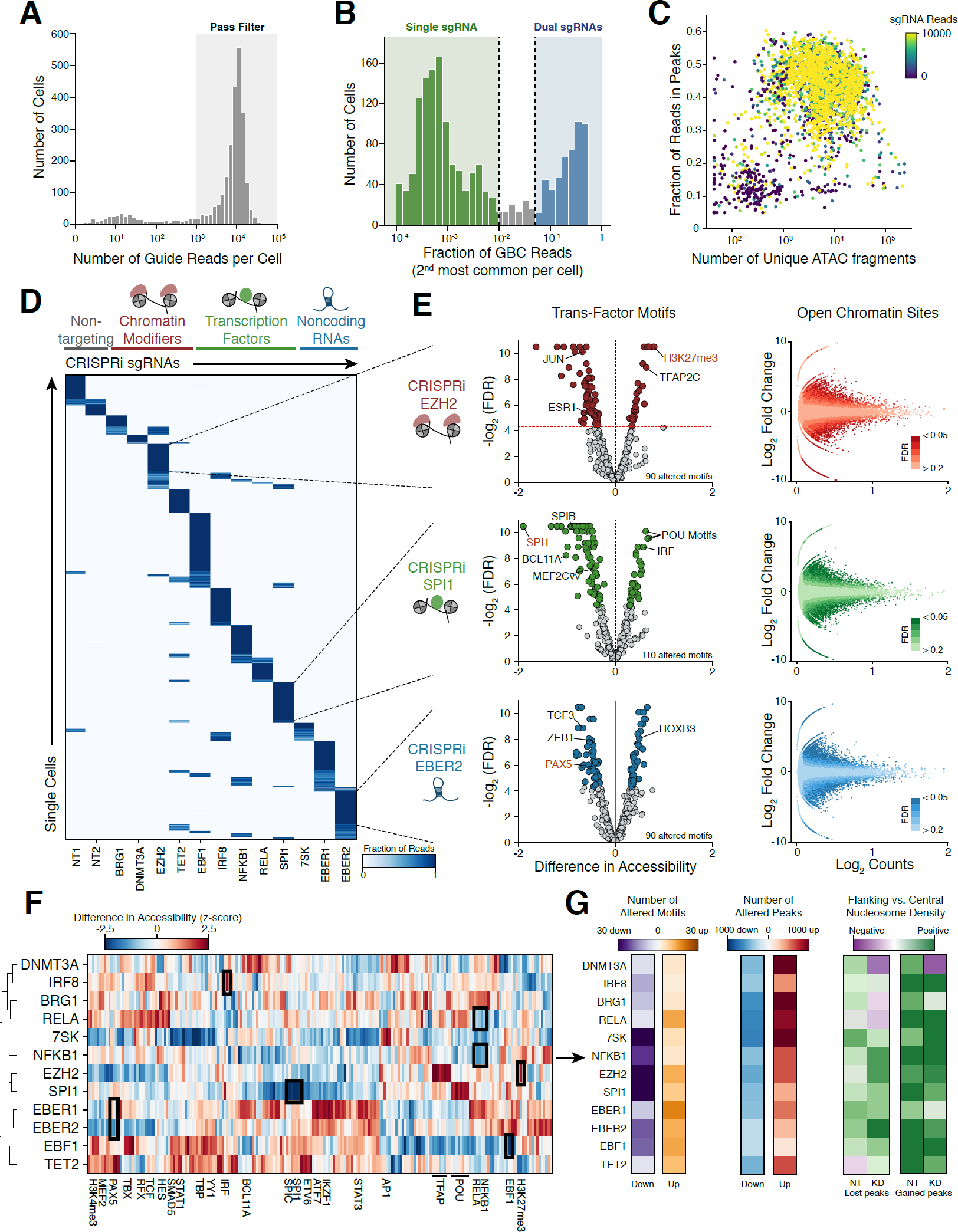
Perturb-ATAC screen for control of accessibility landscape by transcription factors, long non-coding RNAs, and chromatin regulators. **(a)** Histogram of total guide barcode sequencing reads per cell. **(b)** Histogram of the second most common guide barcode identified in each cell. Cells on the low end of the distribution express a single guide RNA, while cells on the high end of the distribution express two guide RNAs. **(c)** Scatter plot of ATAC fragments and fraction of fragments in peak regions. Cells are colored by total guide barcode reads. **(d)** Heatmap of cells (rows) versus guide barcodes (columns) indicating proportion of total reads associated with each barcode. **(e)** Left: volcano plots to identify significantly altered genomic features between cells carrying non-targeting guides and guides targeting EZH2, SPI1, and EBER2 (FDR <= 0.025). Right: Scatter plots of mean accessibility versus fold change of accessibility of individual genomic peaks. **(f)** Heatmap of perturbed factors (rows) versus genomic annotations (columns) indicating difference in accessibility between perturbed cells and non-targeting control cells. Only annotations significantly altered in atleast one perturbation are shown. **(g)** Heatmaps indicating number of significantly altered features (left, absolute log2FC >= 1.5, mean reads/cell >= 0.4), number of altered genomic regions (middle, absolute chromVAR deviation Z >= 0.75, FDR <= .05), or quantification of the ratio of flanking to central nucleosome occupancy at altered peaks (right) for each single perturbation.

We first analyzed ATAC-seq profiles by aggregating single cells based on GBC identity and assessing changes compared to cells receiving NT GBCs (**Figure 3D**). We analyzed chromatin accessibility in the context of three levels of epigenetic regulation: 1) accessibility of *cis*-regulatory DNA elements, 2) TF factor activity genome-wide, and 3) nucleosome positioning at altered peaks. Initial analysis of three regulatory factors confirmed the viability of this approach. For example, depletion of EZH2, a catalytic subunit of the Polycomb repressive complex 2 (PRC2), which deposits repressive H3K27me3 chromatin marks, resulted in significantly increased accessibility at reference regions marked with H3K27me3 (chromVAR accessibility gain 0.78, FDR < .0001; **Figure 3E**). Similarly, the most significant change in TF motif accessibility in *SPI1*-targeted cells was SPI1-motif-containing regions, which showed decreased accessibility in targeted cells (chromVAR accessibility loss 2.17, FDR < .0001; **Figure 3E**). In contrast, SPI-targeted cells showed an increase in accessibility in regions containing IRF motifs, demonstrating that SPI1 controls regulatory factors with activating and repressive functions on chromatin, as has been previously reported (van Riel and Rosenbauer, 2014). Other altered TF motifs in SPI1-targeted cells included BCL11A, SPIB, and MEF2C factors, consistent with the known critical function of SPI1 in regulating factors that are required for B cell lineage commitment (**Figure 3E and Table S1**) (Su et al., 1996; Liu et al., 2003; Stehling-Sun et al., 2009). Finally, targeting of the non-coding RNA EBER2 also identified 90 significantly altered sets of regions, including regions containing the PAX5 motif, a factor that physically interacts with EBER2 to control gene expression (Lee et al., 2015). These results demonstrate that Perturb-ATAC can robustly identify epigenomic phenotypes associated with genetic perturbations of diverse categories of *trans*-factors.

An analysis of all aggregate ATAC-seq profiles derived from 40 single- and double-GBC genotypes revealed accessibility changes in 10,103 open chromatin sites (mean: 404, range: 0-2,250 sites, per genotype) and 833 TF features (mean: 23, range: 0-110 features, per genotype; **Figure 3F, S3A-C, Table S1**). Importantly, cells receiving NT1 or NT2 GBCs showed almost no change in chromatin accessibility compared to all control cells (4 altered open chromatin sites and 0 altered TF features), demonstrating the lack of non-specific epigenomic changes in cells receiving NT sgRNAs (**Figure S3C and Table S1; Methods**). We performed an unbiased clustering of the changes in TF motif accessibility across all perturbations to uncover potential relationships between the perturbed factors (**Figure 3F**). This analysis showed three sub-clusters of *trans*-factors with correlated global epigenomic effects. Cluster 1 factors included the B cell lineage-determining TFs IRF8 and RELA, and the chromatin regulators BRG1 and DNMT3A, suggesting that these factors may function together to establish the open chromatin landscape in B cells (Corces et al., 2016; Lara-Astiaso et al., 2014). Interestingly, two NF?B subunits, NFKB1 and RELA, showed overlapping and distinct effects on chromatin; for example, only RELA perturbation impacted NFYB motif accessibility, while NFKB1 perturbation impacted EBF1 accessibility (**Figures 3F and S3B**). These findings are in line with prior ChIP-seq experiments that demonstrated differences in the binding patterns of NF?B subunits in B cells through the formation of distinct NF?B protein complexes, and highlight the utility of performing high-throughput perturbation screens to identify distinct functions of closely-related factors (Zhao et al., 2014). Similarly, consistent with previous reports of its repressor activity, IRF8 depletion led to increased accessibility at target IRF sites (Tamura et al., 2008). Cluster 2 factors included the small noncoding RNA 7SK, which has been demonstrated to repress transcription of distal enhancers through interactions with chromatin remodeling factors (Flynn et al., 2016), and the repressive chromatin remodeler EZH2 (**Figure 3F**). In addition, cluster 2 factors included NFKB1 and SPI1, which had overlapping effects with repressive factors, as well as factor-specific chromatin activation effects. Cluster 3 included the highly homologous non-coding RNAs, EBER1 and EBER2, as well as TET2 and EBF1. Cells depleted of EBER1 or EBER2 exhibited very similar ATAC-seq profiles, highlighting the functional similarity of these two factors (r = 0.714, p = 2.25e-85; **Figures 3F and S3B**) (Arrand et al., 1989; Samanta et al., 2006), and loss of either factor resulted in altered accessibility at EBF1 sites, consistent with the inclusion of these factors in the same regulatory pathway (**Figure S3B**).

Finally, we used the diversity in ATAC-seq fragment sizes to infer the occupancy and positioning of nucleosomes genome-wide in each perturbation condition (Schep et al., 2015). *Trans*-factors may control accessibility of a locus by regulating the binding of TFs in pre-established nucleosome-free regions as well as by altering the positions or occupancy of local nucleosomes. By analyzing this additional layer of chromatin information, we could assess whether changes in ATAC-seq signal at genomic regions was associated with alterations in nucleosome structure rather than exchange of TF binding within a stable nucleosome scaffold (**Figure S4A**). To validate this approach using Perturb-ATAC data, we examined the profiles of sub-nucleosome-sized and nucleosome-sized fragments surrounding CTCF binding sites in cells expressing NT guide RNAs (**Figure S4B**). As expected, we observed enrichment of sub-nucleosome fragments centered at the CTCF motif, indicative of a central nucleosome-free region. In contrast, nucleosome-sized fragments were enriched in upstream and downstream regions relative to the CTCF motif, representing the well-positioned +1 and -1 nucleosomes previously observed at these sites (Vierstra et al., 2014).

To determine the degree of nucleosome structure dynamics associated with perturbation of each *trans*-factor, we quantified a score representing the flanking accumulation and local depletion of nucleosome-sized ATAC-seq fragments near *trans*-factor binding sites (**Figure 3G**). At regions exhibiting altered total ATAC-seq signal in each perturbation, we compared this nucleosome score between control and perturbed cells. While some factors appeared to operate in the context of stable nucleosome structures, others were associated with substantial alterations in nucleosome profiles. Depletion of either NFKB1 or RELA resulted in a stable nucleosome structure surrounding regions that gain accessibility, suggesting that these factors act as repressors by influencing the binding of other factors in an independently-established nucleosome-free region (**Figure S4C**). DMNT3A may operate differently, however, given that regions that gain accessibility in *DNMT3A*-depleted cells exhibit a stronger central nucleosome structure, consistent with a model in which DMNT3A mediates chromatin repression by active recruitment of negative regulatory factors to an open chromatin region. Reflecting another distinct mechanism, regions losing ATAC-seq signal upon depletion of *7SK* exhibited stronger central nucleosome structure in *7SK*-depleted cells, possibly indicating that 7SK interacts with nucleosome remodeling factors, as has been shown for the BAF complex (**Figure S4C**; Flynn et al., 2016). Overall, these results demonstrate that *trans*-factors regulate nucleosome positioning in distinct ways and that Perturb-ATAC can read out nucleosome structure changes associated with perturbations.

### Discovery of gene regulatory networks controlled by *trans*-factors

We next analyzed Perturb-ATAC profiles to identify epigenomic variability in single cells. We were inspired by previous single-cell analysis frameworks, which demonstrated that natural intercellular heterogeneity could be used to infer regulatory relationships between measured analytes (Klein et al., 2015; Heath et al., 2016). For example, the co-variation of accessibility between sets of regions bound by two distinct *trans*-factors may reflect a common upstream regulator of both factors. In contrast, inverse correlation between two sets of regions may indicate that one factor controls the activity or expression of a negative regulator of the other set of regions. Using this conceptual framework, we could: 1) identify co-varying regulatory networks across single cells, 2) measure the effects of perturbation on each regulatory network, and 3) infer regulatory relationships between the perturbed factor and the constituent factors in the regulatory network (**Figure 4A**).

**Figure 4.**
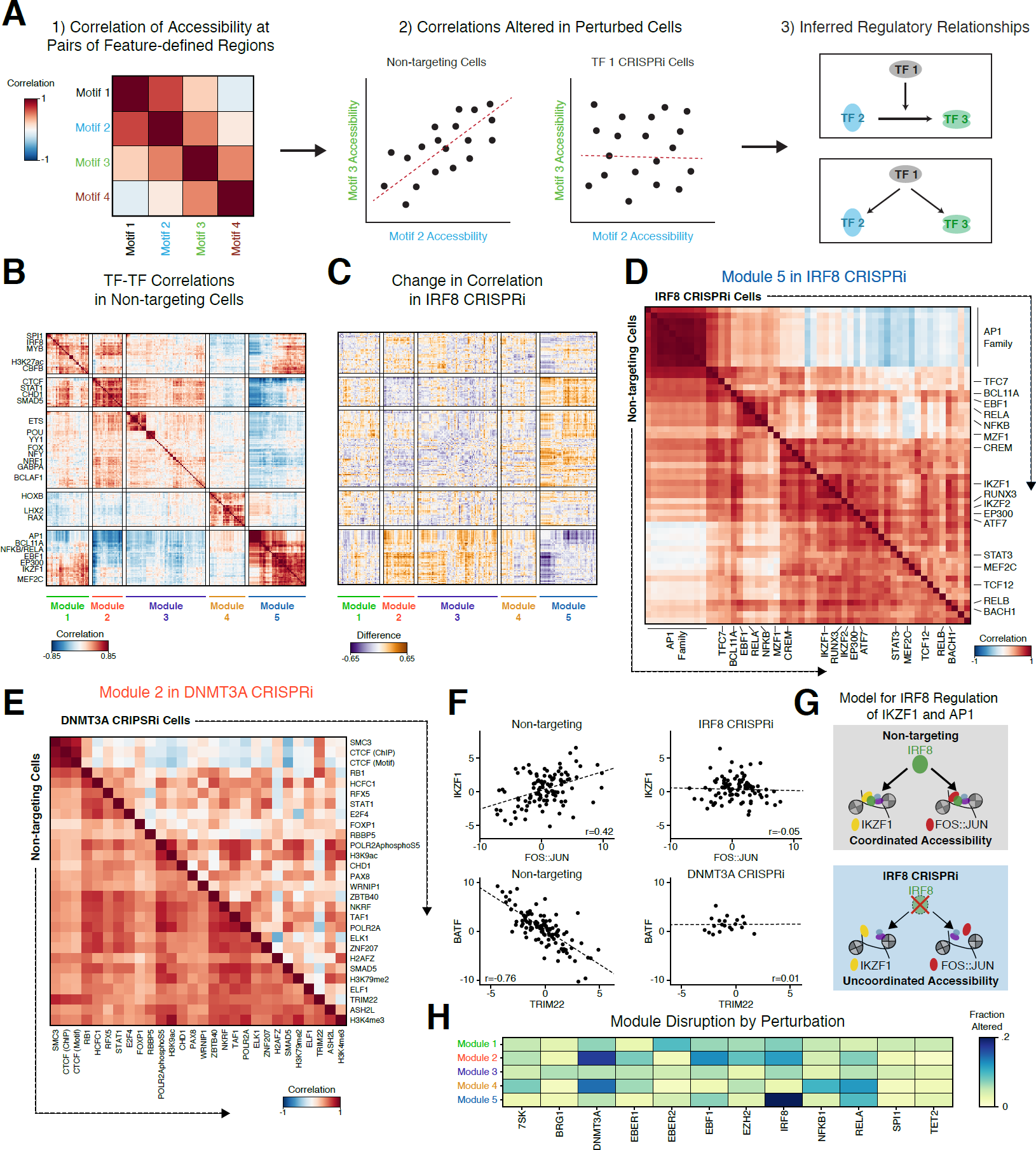
Perturbations influence inter-cellular variability and correlated activity across features. **(a)** Example workflow identifying genomic features exhibiting correlated activity across cells. Left: heat map indicating correlation of motif activity across cells for a group of motifs. Middle: Comparing non-targeting control cells to perturbed cells identifies motif pairs that change in correlation as a result of perturbation. Right: Functional relationships constrain hypothetical regulatory networks. **(b)** Heatmap of Pearson correlations between features across non-targeting cells. **(c)** Heatmap displaying the difference in correlations between non-targeting cells and IRF8 knockdown cells. **(d)** Heatmap displaying Module 5 feature correlations in non-targeting cells (bottom half) and IRF8 (top half) knockdown cells. **(e)** Heatmap displaying Module 2 feature correlations in non-targeting cells (bottom half) and DNMT3A (top half) knockdown cells. **(f)** Scatter plots of accessibility for cells with line of linear best fit demonstrating correlation in specific conditions. **(g)** Hypothetical model of IRF8 co-factor activity with AP1 and IKZF1. **(h)** Heatmap of the fraction of altered feature-feature correlations within modules by perturbation, showing specific effects on particular modules in different perturbations.

We assessed correlations of accessibility across all genomic features (TF motifs or ChIP-seq peaks) in non-targeted cells and identified five modules of correlated feature activity (**Figure 4B and S5A-B; Methods**). Module 1 features, which also correlated with Module 5 activity, were involved in B cell development, including SPI1 and IRF8, as well as general features of active transcription, such as regions exhibiting the active enhancer and promoter chromatin mark H3K27ac. Module 2 included several chromatin regulators, such as CTCF, which controls topological domain structure, and the chromatin remodeler CHD1. ETS factors, which exhibit diverse function in B cell development and immunological function were highly correlated in Module 3 (Nguyen 2012, Blood). This module included YY1, which regulates a distinct subset of chromatin contacts compared to the Module 2 factor CTCF, possibly reflecting exclusive activities. Module 4 contained several homeobox domain TFs and was broadly correlated with a subset of Module 5 features. Finally, Module 5 features primarily included TFs that regulate B cell development, including IKZF1 (also known as Ikaros), BCL11A, EBF1, NFKB, MEF2C and AP1 factors.

Importantly, reconstruction of these regulatory networks in cells depleted of specific *trans*-factors highlighted the subset of modules which were regulated (directly or indirectly) by each perturbed factor (**Figures 4C-E**). For example, depletion of IRF8 led to a substantial re-wiring of the developmental features of Module 5, such that AP-1 factor activity was no longer coordinated with the activity of IKZF1, RUNX, and MEF2C factors, resulting in a decoupling of two normally coordinated modules of developmental TFs (**Figure 4D and S5C**). In particular, IKZF1 and FOS:JUN (AP-1) targets exhibited a strong positive correlation in activity in non-targeted cells (r = 0.420 p = 5.60e-6), which was lost in IRF8 depleted cells (r = -0.051, p=0.627), suggesting that IRF8 coordinates the activity of these two factors (**Figures 4F and 4G**). Indeed, IRF8 has been demonstrated to regulate the expression and activity of IKZF1 in pre-B cells (Ma et al., 2008; Pang et al., 2016). Analysis of regulatory networks in *DNMT3A*-depleted cells revealed changes in the coordinated activity of CTCF with other Module 2 factors, including SMAD5 (**Figure 4E**). Of note, this analysis revealed that the loss of TF-TF co-variation is not dependent on a change in accessibility of each factor, decoupling two distinct mechanisms of TF activity across single cells (**Figures 4E S5D**).

Similar specificity in regulatory interactions could be observed after perturbation of several other TFs and noncoding RNAs (**Figures 4H, S5E-F**). The effect of a perturbation on each module were summarized by counting the number of significantly altered correlations using a permutation-based approach (**Figures S5G and S5H; Methods**). For example, depletion of the B cell developmental regulator EBF1 resulted in substantially altered correlation of Module 2 features, as well as its own Module 5, potentially reflecting the coupled regulation of these two modules in unperturbed cells. The two NF?B subunits, NFKB1 and RELA, exhibited distinct effects on Module 5 features, which include NFKB1 and RELA themselves, while altering Module 4 features to a similar extent (**Figures 4H, S5E**). This relationship extends our observations of both overlapping and specific roles of these two factors in their direct control of chromatin accessibility, possibly associated with their association in two distinct protein complexes. Finally, depletion of EBER2 resulted in altered regulatory interactions between AP-1, IRF, and NFKB factors, suggesting that the expression of virally-encoded RNAs disrupts this regulatory unit, likely through binding and activation of viral sensors (**Figure S5F and Table S1**) (Samanta et al., 2006).

### Mapping epistatic relationships reveals cooperative functions of *trans*-factors in development and disease

To analyze epistatic relationships between *trans*-factors, we generated dual perturbations in single cells for a subset of factors. For each cell, we computed an expected change in chromatin accessibility at each genomic feature based on a multiplicative model of dual, non-interacting perturbations (**Figure 5A; Methods**). We then compared the expected additive result from single-perturbed cells to the observed change in accessibility in dual-perturbed cells and determined the degree of genetic interaction across all genomic features. *Trans*-factor relationships can generally be considered as ‘expected’ based on the combination of the effects of each perturbation alone (additive; suggesting minimal interactions between the perturbations) or ‘unexpected’ (non-additive; suggesting interaction between the perturbations) (Dixit et al., 2016; Jaitin et al., 2016). For example, depletion of EBF1 had no effect on accessibility at SPI1 motif-containing sites, while depletion of SPI1 itself had a strong negative effect on accessibility at these sites (**Figure 5B**). In cells depleted of both EBF1 and SPI1, the observed accessibility matched the expected accessibility based on the combined effect of each factor alone, supporting the notion that these factors do not interact in regulating SPI1 motif-containing sites. Conversely, depletion of either EBER1 or TET2 alone did not affect accessibility at IKZF1 binding sites, yet dual depletion resulted in an unexpected increase in accessibility at these sites (**Figure 5B**), suggesting that these factors converge on a pathway in which each factor alone is dispensable.

**Figure 5.**
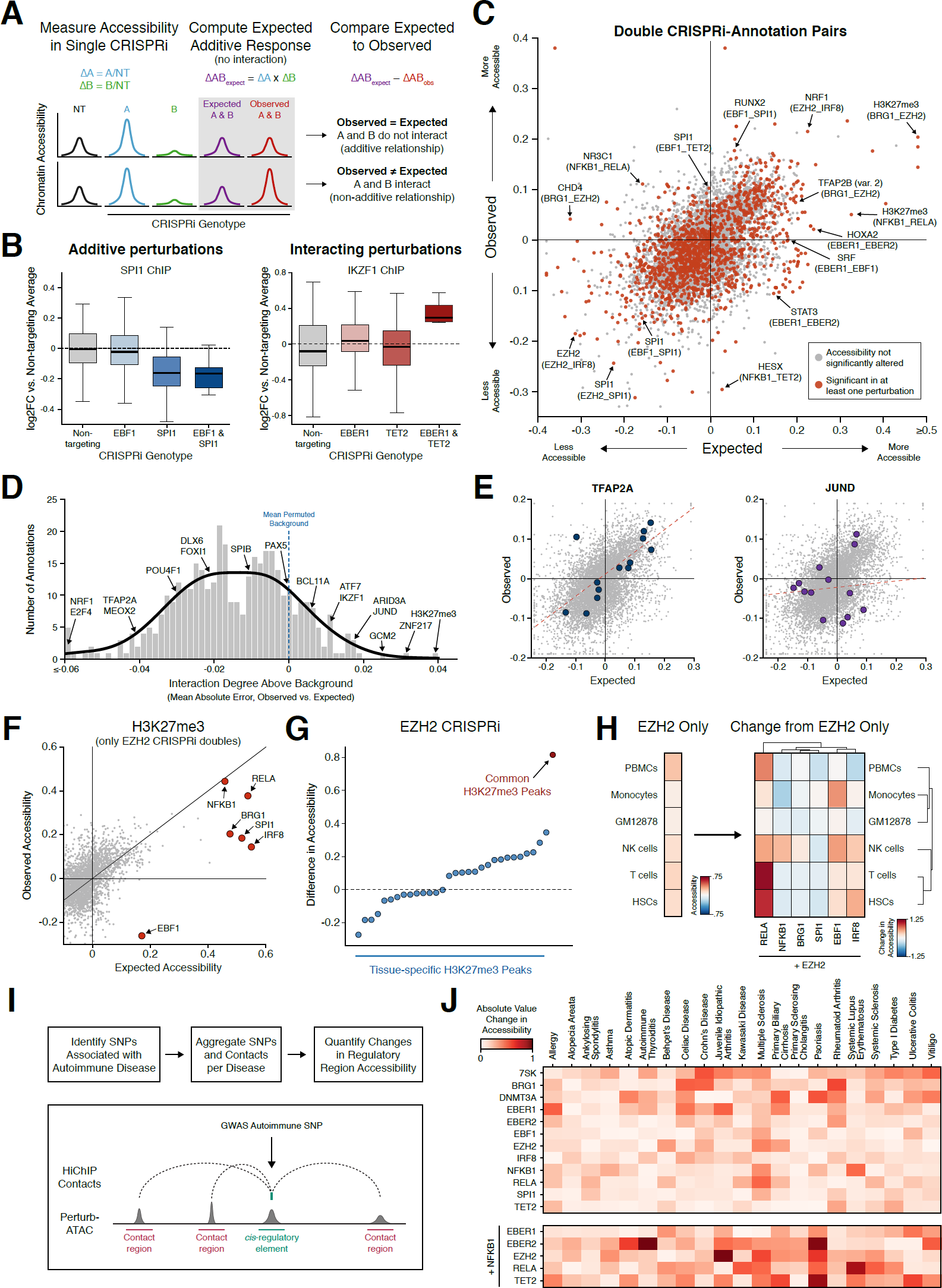
Epistasis analysis identifies functional interaction between a broadly active chromatin regulator and lineage-specific transcription factors. **(a)** Schematic of calculation of expected accessibility in double knockdown context based on additive model integrating accessibility in each single knockdown condition. **(b)** Box plots representing the distribution of SPI1 binding sites (left) and IKZF1 binding sites (right) accessibility for individual cells in respective single or double knockdown conditions. **(c)** Scatter plot of observed versus expected accessibility for epistatic interactions. Each dot represents a single annotation in the pairing of two perturbed factors. Dots highlighted in red indicate significantly altered activity in either single perturbation or double perturbation. **(d)** Histogram of background-corrected interaction degree for each feature. Background distribution calculated by permuting single and double knockdown associations. **(e)** Scatter plots of observed versus expected interactions, highlighting TFAP2A (relatively low interaction degree) and JUND (relatively high interaction degree). **(f)** Scatter plot of observed versus expected change in accessibility at H3K27me3-marked regions in cells depleted of EZH2 and one other factor. **(g)** Scatter plot of relative accessibility in EZH2 knockdown cells compared to control cells for various subsets of H3K27me3 peaks. Common peaks refer to regions exhibiting H3K27me3 status across a majority of cell types. **(h)** Left: heatmap indicating change in accessibility due to EZH2 depletion at regions marked by H3K27me3 in GM12878 and exhibiting H3K27ac mark in each specific other cell type. Right: heatmap indicating change in accessibility in same sets of regions included in the left heatmap, for cells simultaneously depleted of EZH2 and a TF. **(i)** Schematic indicating the workflow to aggregate SNPs associated with autoimmune diseases with 3D chromatin contact regions. **(j)** Heatmap of the absolute change in accessibility for the SNP-contact feature set of each autoimmune disease and perturbation.

We measured the degree of interaction in dual perturbations across all genomic features compared to a permuted background (**Figure 5C and 5D**). Within this analytical framework, features could be categorized according to their regulation by additive or non-additive interactions, revealing features that were generally controlled by independent or synergistic activities of two factors. For example, sets of regions containing TFAP2A motifs were generally regulated by additive interactions (bottom 5% of all features, **Figure 5D and 5E**). In contrast, regions containing JUND motifs were generally regulated by non-additive interactions (top 5% of all features), suggesting synergistic regulation of these regions by the perturbed factors.

Similarly, regions marked by the repressive histone modification H3K27me3 also exhibited a high degree of interaction, in particular, between EZH2 and other *trans*-factors, suggesting a functional relationship between the catalytic subunit of PRC2 and factors which potentially guide PRC2 recruitment or activity at target sites (**Figure 5F**). Since many of interacting factors were B cell lineage-determining factors (EBF1, IRF8, RELA), we reasoned that alternate lineages may be repressed through their combinatorial interactions with EZH2. Indeed, depletion of EZH2 alone primarily resulted in de-repression (increased accessibility) of regions that are commonly marked by H3K27me3 across all cell types and thus may not require additional targeting specificity by other factors, while dual depletion of EZH2 and other factors led to de-repression of regions that are specifically active in alternate hematopoietic lineages (**Figures 5G and 5H**). Moreover, each factor analyzed by dual depletion with EZH2 resulted in a unique set of cell type-specific de-repressed regions. For example, EBF1 cooperated with EZH2 to repress alternate monocyte and natural killer cell fate, while IRF8 and RELA cooperated with EZH2 to repress progenitor fates, perhaps reflecting their function in the step-wise development of B cells (**Figure 5H**).

Finally, we asked whether Perturb-ATAC analysis could inform regulators of noncoding regions that contain genetic variants associated with human disease. We examined this in the context of 21 autoimmune diseases, a subset of which demonstrate an enrichment of causal variants in B cell-specific enhancers (Farh et al., 2015). We generated a feature set for each disease that consisted of all ATAC-seq peaks within 10kb genomic windows of causal variants and their 3D enhancer-promoter connections (using GM12878 H3K27ac HiChIP) (Farh et al., 2015; Mumbach et al., 2017). We then measured the effect of each perturbation, in isolation or combination with other factors, on the accessibility of each disease feature set (**Figure 5I**). Strikingly, this analysis revealed disease-specific activities of several TFs, which were supported by previous binding-site predictions (**Figure 5J**). For example, NFKB binding sites, and B cell enhancers generally, were previously shown to be highly enriched near causal variants associated with multiple sclerosis (MS) and system lupus erythematosus (SLE) (Farh et al., 2015), and these sites demonstrated changes in accessibility in single cells where *NFKB1* or *RELA* were depleted (**Figure 5J**). In contrast, although variants associated with celiac disease (CD) and rheumatoid arthritis (RA) are also enriched in B cell enhancers, these diseases showed a distinct perturbation signature. CD and RA variant-associated enhancers were most strongly impacted by perturbation of chromatin modifiers BRG1 and DNMT3A (**Figure 5J**). Moreover, analysis of cells that received dual perturbations revealed variant-associated changes that could not be observed with single perturbations. For example, although perturbation of NFKB1 did not alter many variant-enhancers in isolation, dual perturbation of NFKB1 and several other factors revealed interactions that resulted in accessibility change at peaks associated with several diseases (**Figure 5J**). Taken together, these results demonstrate that epistatic interactions represent a layer of epigenomic control in development and disease that cannot be simply predicted by analysis of individual perturbations, and that Perturb-ATAC presents a scalable platform for discovering and mapping such interactions.

### The regulatory landscape of human epidermal differentiation

Dynamic biological systems such as tissue differentiation present a unique opportunity for pooled, high-content screens to assess the function of many *trans*-factors while internally controlling for experimental variation. The human epidermis is a constantly renewing, stratified epithelial tissue that protects the body from environmental insults and provides structure for hair follicles, sweat glands, and pigmenting melanocytes (Gonzales et al., 2017). The process of terminal epidermal differentiation involves dynamic expression of thousands of genes as progenitor cells migrate from the basal layer, begin keratinization, and ultimately undergo cornification to form the outermost protective layer of the skin (Lopez-Pajares et al., 2015). We sought to understand how transcription factor control of chromatin state regulates this process by perturbing key TFs during differentiation.

We first assessed the landscape of chromatin accessibility in normal keratinocyte differentiation to identify key regulators of this process as candidates for Perturb-ATAC. We employed a model of epidermal differentiation in which human primary progenitor keratinocytes are either cultured to mimic the basal progenitor population in tissue or seeded at high density with high calcium to trigger the differentiation process (**Figure 6A**) (Kretz et al., 2013). We performed single-cell ATAC-seq on populations of cells from each of three time points in this model to capture undifferentiated cells (Day 0), mid-differentiation (Day 3), or late differentiation (Day 6). Based on the accessibility of genomic features, single cells largely separated in reduced dimensionality space based on their differentiation time point (**Figure 6B**). However, a small number of “precocious” differentiating cells were observed in the Day 0 population, and the Day 6 population separated into one Day 6-specific subset and a subset that appeared more similar to Day 3 cells, representing endogenous heterogeneity in each population observable only using single-cell analysis. Placement of each cell along a one-dimensional pseudotime trajectory highlighted the continuous nature of the differentiation process (**Figure 6B**) (Qiu et al., 2017).

**Figure 6.**
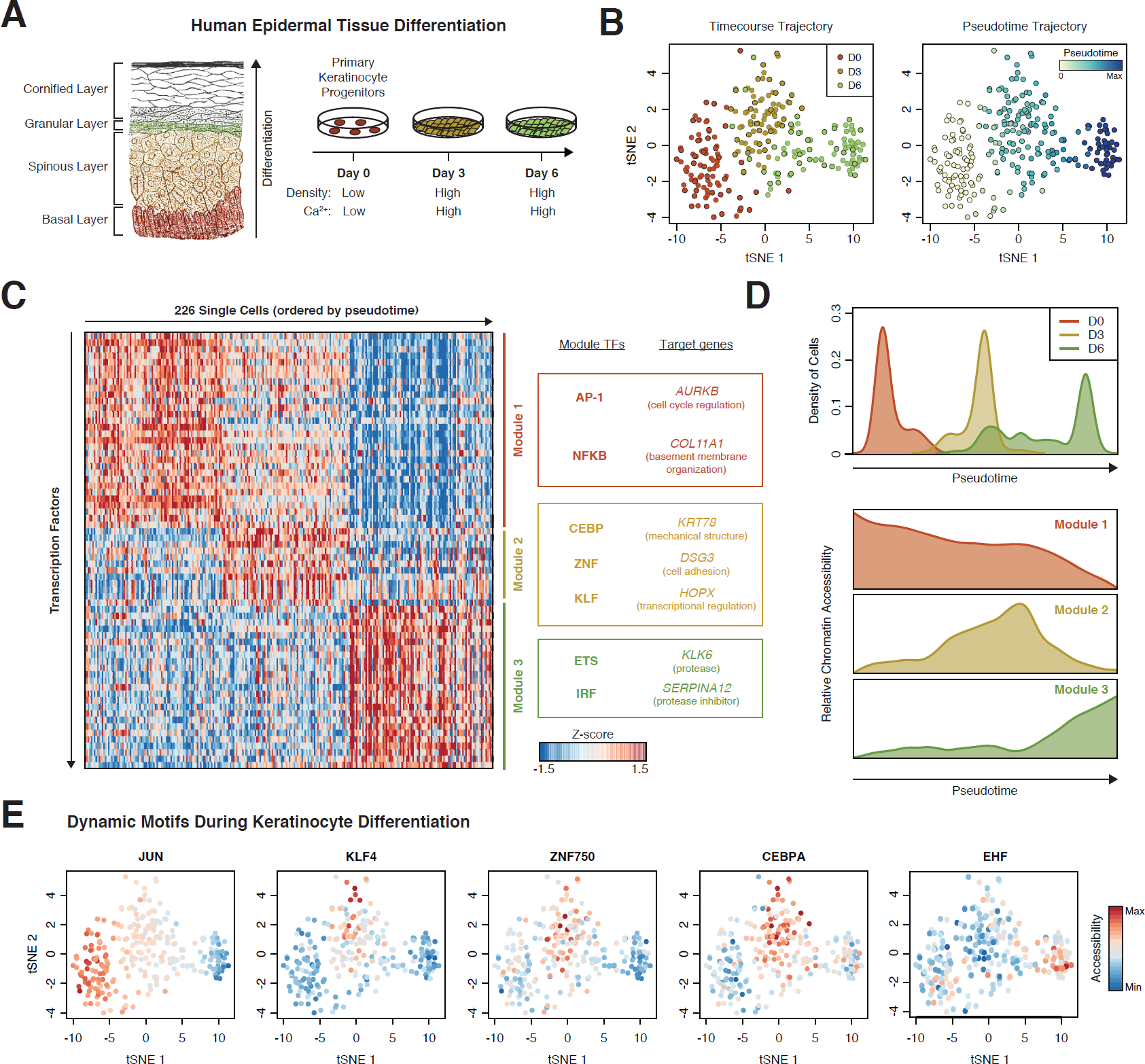
Modules of transcription factors exhibit distinct temporal activity in epidermal differentiation. **(a)** Schematic of human epidermis and cell culture model system of epidermal differentiation. **(b)** tSNE projection of TF feature activity for epidermal cells labeled by differentiation day (left) or pseudotime (right). **(c)** Heatmap of cells ordered by pseudotime (columns) versus TF feature activity (filtered for motifs with dynamic activity). Modules represent collections of TF features with similar temporal profiles. Genes listed next to heat map were found proximal to (<50kb) from genomic regions exhibiting accessibility kinetics associated with that module. **(d)** Top: density histogram of pseudotime values for cells from each day of differentiation. Bottom: Average accessibility profiles for each module identified in (c). **(e)** tSNE projections of TF activity, cells are labeled by relative activity of individual motifs for each plot.

We next determined a set of *trans*-factors to perturb by identifying highly dynamic *cis*-regulatory features during keratinocyte differentiation and the *trans*-factor binding sites enriched in these regulatory elements. We analyzed the chromatin accessibility of 94,633 *cis*-elements at the level of 411 transcription factor motifs and ChIP-seq peaks (Schep et al., 2017). By comparing the chromatin accessibility of each set of TF target regions in each cell with the position of that cell on the differentiation pseudotime, we identified 67 TFs whose chromatin dynamics suggested a role in regulating keratinocyte differentiation (**Figure 6C**). These TFs clustered into three modules of dynamic behavior (**Figures 6C and 6D**). The first module (30 features, including AP-1 and NFKB target sites) exhibited relatively high accessibility in progenitor cells and a steady decrease in accessibility as differentiation progressed. Genes proximal to specific regions following Module 1 kinetics included *AURKB*, a regulator of mitosis, and *COL11A1*, a component of the basement membrane. The behavior of these elements is consistent with roles for these genes in progenitor cell functions that are lost as cells detach from the basement membrane and initiate terminal differentiation. Module 2 (12 features, including target regions of CEBP and KLF family factors, as well as the TF ZNF750), exhibited high accessibility specifically during mid-differentiation. The selective activity of these factors midway through differentiation was not previously seen in bulk experiments, likely due the presence of mixed populations of mid- and late-differentiated cells observed in the Day 6 culture (Rubin et al., 2017) (**Figure 6B)**. The genes *KRT78, DSG3*, and *HOPX* were proximal to regions exhibiting Module 2 dynamics, representing the distinct programs of keratinization, cell-cell adhesion, and transcriptional regulation, respectively, engaged by mid-differentiating cells. Finally, Module 3 (25 features) included target regions of ETS and IRF family factors, and specifically gained accessibility in late differentiation. Associated genes included *KLK6* and *SERPINA12*, a protease and protease-inhibitor, respectively, which encode extracellular proteins that regulate the coordinated release (desquamation) of cells from the outer layer of the epidermis (Guttormsen et al., 2008; Kishibe et al., 2007).

We used this analysis to prioritize a set of candidate TFs for characterization with Perturb-ATAC in order to understand their precise roles in governing the chromatin state dynamics of differentiation. Notably, many of the TFs associated with each module have reported roles in the epidermis, supporting the ability of scATAC-seq data to identify key factors associated with a dynamic process. For a candidate regulator of AP1 motif-containing regions, which are associated with Module 1, we selected JUNB, an AP1 family member previously shown to exert control over epidermal homeostasis and tumorigenesis (Eckert et al., 2013). We reasoned that as Module 2 features gain activity during keratinocyte differentiation, they might be critical to the initiation of differentiation and thus chose to perturb three regulators: 1) KLF4, a factor that promotes epidermal differentiation gene induction, 2) ZNF750, a factor reported to engage in positive and negative gene regulation in epidermal differentiation, and 3) CEBPA, which has known role in murine epidermal homeostasis (Lopez et al., 2009; Sen et al., 2012). As a Module 3 regulator, we chose EHF, an ETS family member which is selectively expressed in epidermal cells and has been implicated in controlling epidermal differentiation (Rubin et al., 2017). Closer analysis of these TFs confirmed patterns of dynamic accessibility matching their parent module (**Figure 6E**). Importantly, by leveraging the Perturb-ATAC platform, we aimed to not only study the individual activities of these TFs but also to comprehensively map their pairwise interactions.

### Perturb-ATAC screen for TF control of cellular differentiation trajectories

For the analysis of keratinocyte differentiation, we developed a secondary version of the Perturb-ATAC protocol in order to achieve two goals: 1) to directly detect the sgRNA itself rather than a GBC, and 2) to assess the effects of CRISPR gene knockouts rather than CRISPR interference. This alteration was motivated by the desire to make the Perturb-ATAC protocol directly amenable to standard CRISPR knockout sgRNA libraries, rather than limiting its use to GBC-containing CRISPR constructs. In this experimental workflow, primary keratinocytes were first transduced with a lentivirus encoding Cas9 and selected for successful transduction. Subsequently, Cas9-expressing keratinocytes were transduced with either one or two sgRNA cassettes, corresponding to the five TFs (JUNB, KLF4, ZNF750, CEBPA, EHF) or two NT control sgRNAs, producing both single- and double-targeted cells (**Figures S6A and S6B)**. To allow for Cas9 activity and genomic disruption of the targeted genes, keratinocytes were maintained in undifferentiated culture conditions for 7-10 days. Pooled cells were then transferred to differentiation culture conditions and harvested at Day 3 for Perturb-ATAC analysis.

We implemented a workflow to directly detect sgRNA sequences by modifying the previously described protocol for detecting GBCs. To detect sgRNAs, we performed reverse transcription using a reverse primer that matched the common 3’ end of the sgRNA, followed by subsequent PCR amplification using a pool of forward primers matching the variable 5’ ends of the sgRNAs used in the experiment (**Figures 7A and S7A)**. A similar analytical framework as described previously was used to analyze sgRNA sequencing reads and assign Perturb-ATAC genotypes to individual cells. First, we set a plate-specific depth cutoff to distinguish successful and failed sequencing reactions (**Figure S7B**). We found that an average of 87.9% of reads mapped to an sgRNA sequence, indicating a low level of off-target amplification (**Figure S7C**). Cells were then analyzed to determine the proportion of reads attributed to the expected background sgRNAs, allowing us to exclude cells exhibiting high background reads (cells with greater than 1% background reads were excluded from analysis) (**Figure S7D**). The remaining cells were then assessed to determine the proportion of reads attributed to the first and second most common sgRNA, resulting in a bimodal distribution reflecting the expected presence of one or two distinct sgRNAs in each cell (**Figure S7E**). Cells ultimately designated as single or double sgRNA-expressing exhibited clear separation into distinct populations (**Figure S7F**). Altogether, we identified 279 single sgRNA-expressing cells and 235 double sgRNA-expressing cells, encompassing 23 distinct genotypes of all individual and double CRISPR perturbations as expected from our transduction protocol (**Figure 7B**).

**Figure 7.**
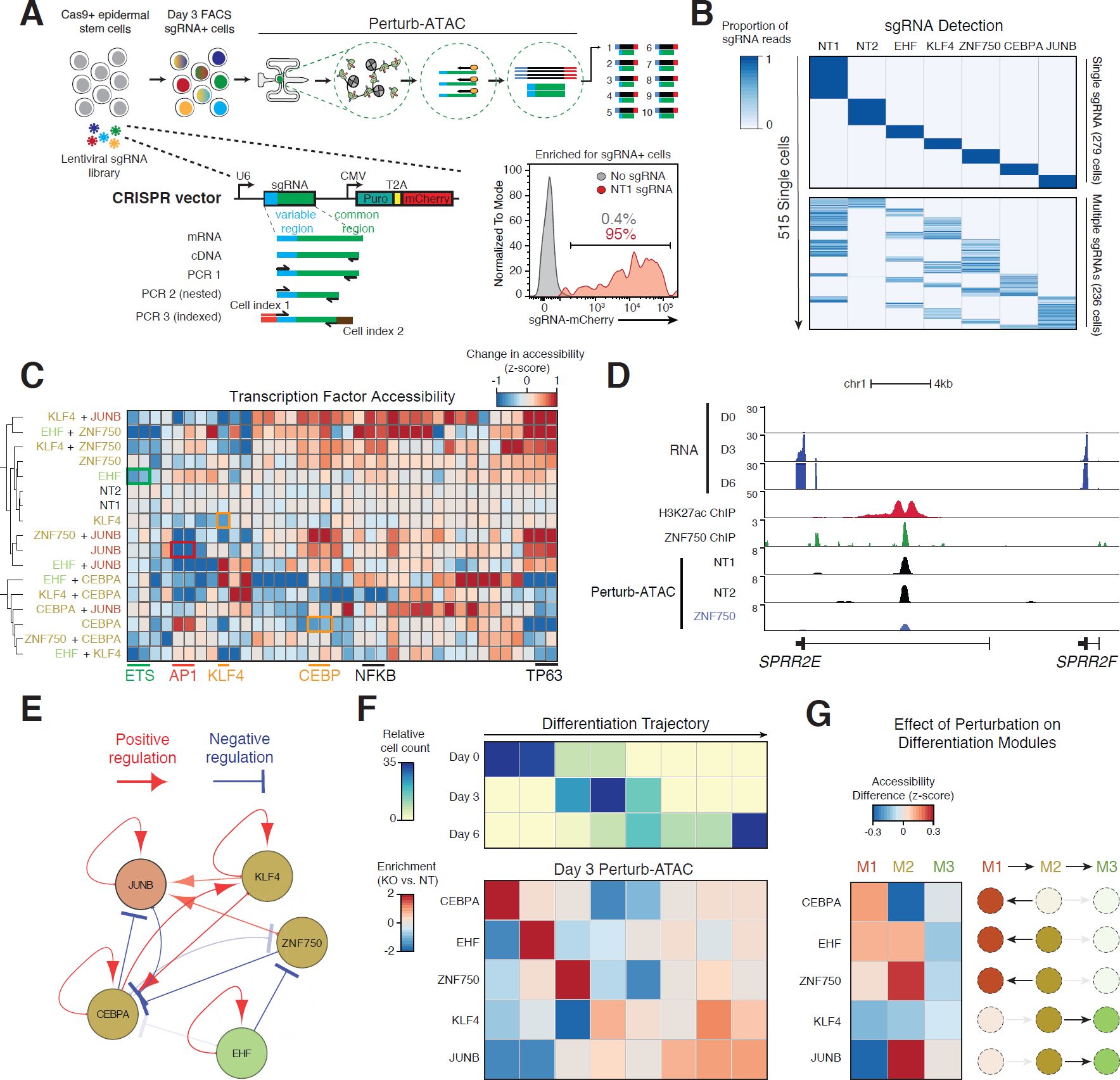
Multiplex knockout screen of transcription factors in differentiation. **(a)** Schematic of sgRNA expression vector and library amplification for direct sequencing readout of guide RNA identity. **(b)** Heatmap of sgRNA identities (columns) versus single cells (rows) indicating the proportion of all reads associated with each sgRNA. **(c)** Heatmap of genetic perturbations (noted by target gene) versus TF features indicating activity of TF feature in perturbed cells relative to non-targeting (NT) cells. Similar motif features from AP1, FOX, and ETS families were merged. **(d)** Genomic locus of SPRR2E gene. Perturb-ATAC tracks represent signal from merged single cells identified for each sgRNA. H3K27ac and ZNF750 ChIP-seq tracks (from Day 3 differentiating keratinocytes, normalized to 10m reads, from Rubin et al. 2017). **(e)** Representation of positive and negative regulation between targeted genes (factors) and sets of genomic regions (features). Arrows are shown for regulation with FDR < 0.25 and decreasing transparency is associated with lower FDR. Map was generated using Cytoscape v3.1.0. **(f)** Top: heatmap displaying the frequency of cells in each of eight bins representing progression along differentiation trajectory. Bottom: heatmap indicating the enrichment or depletion of cells in each differentiation bin compared to non-targeting control cells. For each perturbation, a custom reduced dimensionality space was created to highlight altered features. **(g)** Heatmap of perturbations (targeted genes, rows) versus modules of features (columns). For each module, the mean change in feature activity is shown.

We identified 399 features altered across all TF knockout cells (FDR < 0.1). These features included multiple instances of TFs regulating accessibility of their target sites, as expected: regions containing motifs or ChIP-seq peaks corresponding to JUNB, EHF, KLF4, and CEBPA all exhibited reduced accessibility upon depletion of each respective factor (FDR < 0.05) (**Figure 7C**). Notably, ZNF750 target regions were not uniformly more or less accessible in *ZNF750*-targeted cells, consistent with the previously reported role of this factor as a positive and negative gene regulator (Boxer et al., 2014). For example, the intronic enhancer of the epidermal cornified envelope gene *SPRR2E* is bound by ZNF750 and exhibited loss of accessibility upon *ZNF750* disruption, while other ZNF750 binding sites gained accessibility upon knockout (**Figure 7D and S8A**). Globally, *ZNF750* knockout resulted in a nearly even total of 649 gained and 620 lost peaks (FDR < 0.01, FC>1). Surprisingly, Module 2 factors, which exhibited the highest level of accessibility at mid-differentiation, displayed diverging roles in keratinocyte differentiation. For example, perturbation of *ZNF750* led to increased accessibility at TP63 and NFKB motif sites, which are two factors known coordinately regulate epidermal identity and progenitor maintenance (Yang et al., 2011). In contrast, perturbation of *CEBPA* led to decreased accessibility of TP63 and NFKB features. This unexpected divergence in the behavior of Module 2 factors highlights the importance of unbiased perturbation and phenotyping to uncover distinct *trans*-factor activities that occur as cells progress in differentiation. Moreover, a global analysis of the perturbed factors and their corresponding target regions uncovered a highly inter-connected network of regulation; namely, every perturbed factor altering the accessibility of at least one other factor’s target regions (**Figure 7E**).

To comprehensively assess the effects of TF perturbation on the trajectory of keratinocyte differentiation, we projected perturbed cells onto a pseudotime axis derived from wildtype Day 0, Day 3, and Day 6 cells. For each perturbation, we assessed the distribution of individual cells along the differentiation pseudotime to identify heterogeneity in cell states induced by TF perturbations (**Figures 7F, S8B and S8C**). We also determined the average change in accessibility for features in each of the three differentiation regulatory modules identified earlier in unperturbed cells (**Figure 7G**). This analysis revealed a critical role for CEBPA in initiating the regulatory program of keratinocyte differentiation, as *CEBPA* knockout cells were enriched at the early differentiation stage but depleted at the expected mid-differentiation state. Correspondingly, *CEBPA* knockout cells displayed a decrease in accessibility of Module 2 features but increased accessibility of Module 1 features, indicating that *CEBPA* knockout cells remained in a progenitor state and were unable to fully engage the mid-differentiation program. In contrast, *KLF4* knockout cells were depleted at early differentiation stages but enriched at later differentiation, again highlighting the divergent roles of *trans*-factors in regulating mid-differentiation: *CEBPA* knockout caused cells to move back to an undifferentiated state, while *KLF4* knockout caused cells to progress along differentiation to later states.

Cells depleted of the Module 1 factor JUNB were enriched in mid-late differentiation stages and strongly depleted in the early differentiation state. These cells had decreased accessibility of Module 1 factors but increased accessibility of Module 2 and 3 factors (**Figure 7G**). This result is consistent with a role for JUNB in maintaining cells in a progenitor state, where its activity is highest in unperturbed differentiation, and preventing their progression to a differentiated state. Together, these results demonstrate the power of a single Perturb-ATAC experiment to map the roles of *trans*-factors in tissue differentiation. By analyzing perturbations from a pooled experiment with single-cell resolution, we identify both the specific regulatory factors responsible for distinct cell states as well as rerouted differentiation trajectories that occur in response to TF perturbation in the human epidermis.

### Comprehensive epistatic mapping reveals the logic of regulatory synergy

Since all possible dual perturbations were included in the epidermal differentiation Perturb-ATAC experiments, we were able to extend our previous findings on TF epistasis. As described previously, we calculated an expected multiplicative effect of dual factor depletion based on the effect of every single perturbation on each genomic feature. We then scored each perturbation-feature pair as additive (no interaction; observed similar to expected), synergistic (positive interaction; observed greater than expected), or buffering (negative interaction; observed less than expected) (**Figures 8A and 8B, S8D**). This workflow identified 344 additive relationships, 102 synergistic interactions, and 101 buffering interactions across all pairwise perturbations. Unlike in other systems, we observed only a weak correlation between the magnitude of a single perturbation on genomic features and the degree of genetic interaction for that feature (Costanzo et al., 2010) (**Figure S8E**).

**Figure 8.**
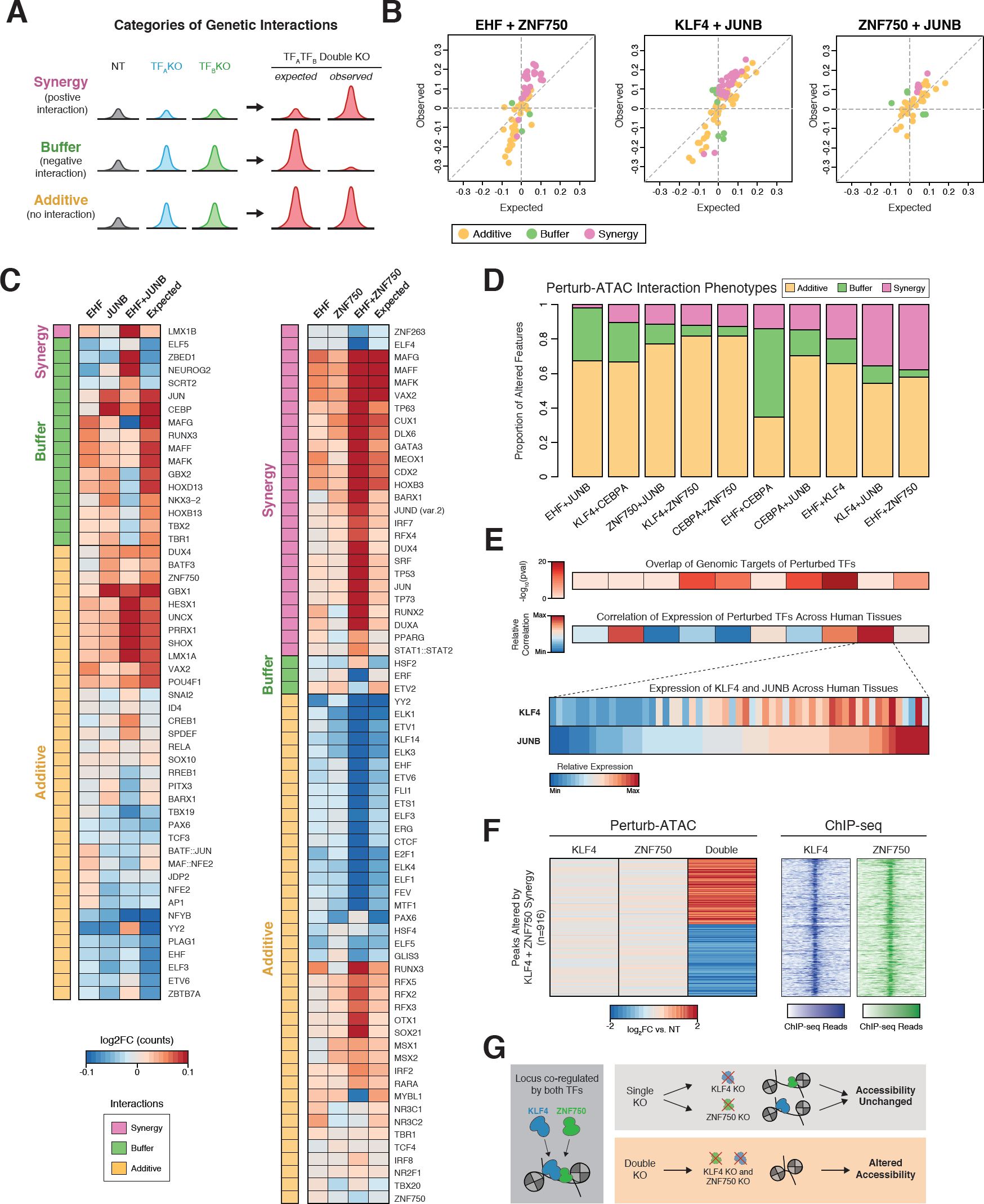
Pairs of perturbations exhibit distinct patterns of epistatic interactions. **(a)** Example representative peak signal for each category of interaction. **(b)** Scatter plots of observed versus expected (based on additive model) accessibility in double knockout cells. Only features significantly altered in either single knockout or double knockout condition are plotted, and feature colors indicate category of interaction. **(c)** Left: heatmap of altered activity of features (rows) in the condition of EHF knockout, JUNB knockout, or simultaneous EHF and JUNB knockouts in the same cell, along with their expected activity. Right: Similar to left, for EHF and ZNF750 knockouts. **(d)** Bar plot of the proportion of interacting features belonging to each category. Each column represents a particular pair of targeted genes. Only features altered in either single perturbation or the double perturbation condition are considered. **(e)** Top: Heatmaps indicating significance of genomic overlap or correlation of gene expression for pairs of TFs corresponding to pairs displayed in (d). Bottom: Heatmap displaying relative RNA expression of KLF4 and JUNB across tissues from the Roadmap Epigenomics Project. **(f)** Left: Heatmap indicating relative accessibility of genomic regions (rows) exhibiting synergistic behavior in KLF4 and ZNF750 double knockout cells. Right: heatmap with rows corresponding to regions displayed on left, displaying ChIP-seq profiles for KLF4 and ZNF750. **(g)** Hypothetical model of KLF4 and ZNF750 redundancy for maintenance of accessibility at co-occupied loci.

The investigation of particular pairs of factors knocked-out simultaneously revealed surprising interactions of previously identified relationships. For instance, single knockout of *JUNB* led to an increase in accessibility of CEBP motif-containing sites while the dual *EHF* and *JUNB* knockout had a much more modest effect, reflecting a negative interaction between EHF (which itself has a weak effect on CEBP sites) and JUNB (**Figure 8C**). This result may reflect a dependence between Module 2 activities (including CEBP) and Module 3 TFs (including EHF). However, JUNB and EHF had no evidence of interaction at their respective target sites, suggesting an indirect effect mediated by these TFs serving as co-factors.

When comparing all pairs of targeted TFs, we noticed large disparities in the prevalence of each interaction category (**Figure 8D**). By quantifying the proportion of interactions exhibiting additive, buffering, or synergistic behavior, we observed over a three-fold difference in the prevalence of synergistic interactions across TF pairs. EHF and JUNB had a relatively low degree of synergy, in contrast to the high degree of synergy observed for EHF and ZNF750. This synergy was particularly apparent at TP63 family motifs where the TF TP63 binds to exert its function as a master regulator of epidermal homeostasis and differentiation (Bao et al., 2015). While *EHF* and *ZNF750* knockout alone exhibited modest effects on TP63 motif sites, dual knockout markedly increased their accessibility, potentially indicating that these factors antagonize the activity of the TP63-interacting chromatin remodeler BRG1 (**Figure 8C**).

Synergy between two factors may be a consequence of perturbed factors regulating a pathway in similar ways. We reasoned that a potential mechanism to explain synergy between two factors’ regulation of chromatin state could be the genomic co-localization of those factors, analogous to the genetic interactions due to physical protein interactions observed in yeast (Costanzo et al., 2010). For example, if a given genomic element is separately bound by two *trans*-factors, which each positively regulate its accessibility, then each factor alone may be dispensable for chromatin accessibility at that element. However, simultaneous disruption of both factors would lead to failed recruitment of other machinery required to maintain the chromatin state of that element, manifesting as altered accessibility observed in double knockout cells, but masked in single knockouts. To test this hypothesis, we computed the degree of genomic overlap for the target regions of each pair of perturbed factors (**Figure 8E**). Indeed, the factor pairs with the highest prevalence of synergistic interactions exhibited greater overlap of target sites, consistent with the co-binding model (Spearman’s correlation = 0.4). Beyond co-binding, we reasoned that coordinated patterns of gene expression across different cell states could indicate that two factors act in similar pathways and thus may functionally interact. To examine the relationship between co-expression and epistatic synergy, we calculated the correlation of expression between all pairs of TFs studied across a diverse set of tissues. Interestingly, the correlation of expression across tissues was associated with the level of synergy (Spearman’s correlation = 0.358). Co-expression across tissues was exemplified by KLF4 and JUNB, which exhibit the second greatest degree of synergy across pairs and correlated expression (**Figure 8E**). This result may reflect the need for strict regulation of cooperative TF expression across tissues in order to achieve distinct cell states.

To further explore the relationship between genomic co-binding and synergy suggested by our data, we closely examined the KLF4 and ZNF750 pair, which exhibited a high degree target-region overlap. We asked whether the specific genomic regions at which synergy was observed (displaying a greater change in double than single perturbations) were co-bound by both factors, in accordance with the model where overlap of TF target-regions drives epistatic synergy. Using previously published ChIP-seq data for KLF4 and ZNF750 (Boxer et al., 2014), we observed that these regions (916 loci) were in fact commonly bound by both factors, providing strong support for the model that *trans*-factor co-binding underlies coordinated regulation of shared loci (**Figures 8F and 8G**).

We interpret these results as a mechanism of epigenetic robustness, whereby cells rely on functional redundancy to buffer perturbations of regulatory factors. Importantly, we note that this observation was made possible by the inclusion of both single and double knockouts in our Perturb-ATAC experiment, which was enabled by the highly scalable nature of the platform to many genetic perturbations. By comprehensively mapping the epistatic relationships in a model of human epidermal differentiation, we uncover specific examples of functional genetic interactions. More broadly, we reveal genomic co-localization as a cellular mechanism to ensure epigenetic robustness, allowing cells to tolerate individual variations in *trans*-factor expression but maintain key regulatory processes of chromatin state in dynamic systems.

## Discussion

In this study, we developed Perturb-ATAC, which bridges the gap between technologies that perform high-throughput genetic screens and technologies that measure chromatin state genome-wide. Until recently, high-throughput genetic screens have largely relied on the introduction of genetic variants in a population of cells (for example, using CRISPR/Cas9), followed by the selection of cells containing variants that confer a limited set of cellular phenotypes, such as increased cell growth or viability. However, identifying genetic variants that regulate genome-wide chromatin states, which are challenging to identify with standard selection protocols, requires focused experiments, where candidate factors are perturbed and then tested one-by-one. This experimental bottleneck has dramatically limited the scale of factors and their interactive impacts on chromatin states that can be examined in any given experiment. Therefore, understanding factors that regulate chromatin state remains a significant challenge.

Recently, several studies described the coupling of CRISPR screens and single-cell RNA sequencing, which enables screening for high complexity phenotypes using single-cell genomics. There are two overarching benefits of such an approach: 1) pooled cells bearing distinct perturbations, as well as control cells, can be grown in precisely the same conditions, minimizing the impact of experimental variation on the measured phenotype, and 2) experiments can be performed at an otherwise inaccessible scale, since a far greater number of perturbations than could be easily measured one-by-one can be simultaneously assayed in a single experiment. We have applied this concept further to now enable the coupling of CRISPR screening and readouts of epigenomic state in single cells. By achieving simultaneous measurement of the perturbation identity and ATAC-seq profiles in individual cells, we provide an easily-scalable platform for discovery of complex regulatory interactions. We applied this method to determine the roles of a diverse set of *trans*-regulatory factors, including transcription factors, chromatin modifiers, and human and viral noncoding RNAs. Perturb-ATAC data enables the study of several layers of chromatin regulation: 1) individual *cis*-regulatory elements, 2) inferred TF activity from *cis*-regulatory modules, and 3) nucleosome positioning and occupancy. Thus, Perturb-ATAC may be particularly well-suited for the examination of the precise molecular switches in the non-protein-coding genome that underlie cell states, compared to existing methods that pair whole transcriptome profiles with perturbations in single cells.

The single-cell resolution of this assay allowed us to perform several rounds of data analysis to understand the interplay of regulatory factors that underlie chromatin state. First, we used the data to perform ‘pseudo-bulk’ experiments, whereby single-cells were grouped according to their perturbation and treated as an aggregate profile to identify changes in accessibility genome-wide compared to control cells. Indeed, this analysis closely mirrored the results that were obtained from experiments done in bulk cell populations, providing a high-throughput platform to assay epigenomes in perturbed cells. However, we could further exploit the naturally-existing single-cell variation within a single perturbation to identify the correlated activity of TFs that form ‘regulatory modules.’ By examining regulatory modules across perturbations, we could infer functional relationships between TFs. For example, in B cells, we identified a TF module consisting of several known B cell lineage factors, whose activity was coordinated by IRF8. In the absence of IRF8, the loss of coordinated activity within this module led to the altered accessibility of approximately 1,000 *cis*-elements. Finally, we could perform a third layer of analysis by comparing cells that received multiple perturbations to those that received only single perturbations to map epistatic interactions between factors. This analysis uncovered a surprising degree of interactions that were ‘hidden’ in the analysis of only single perturbations. For example, in epidermal cells, the epistatic activity of KLF4 and ZNF750, driven by their co-occupancy at *cis*-elements, was crucial to the terminal differentiation of keratinocytes, consistent with prior studies in epidermis (Boxer et al., 2014). Thus, multiplexed Perturb-ATAC screening offers the ability to uncover new layers of gene regulation that may not be visible in bulk single perturbation experiments.

Importantly, these molecular insights can be integrated into the overall cellular trajectories of differentiation that can be constructed from single-cell epigenomic data. Previous studies identified the presence of significant intra-population variability in the setting of hematopoiesis and embryogenesis and highlighted regulatory insights – such as epigenetic regulators of rare cell types or intermediate progenitors – that could only be observed with single-cell analysis (Buenrostro et al., 2017; Cusanovich et al., 2018; Satpathy et al., 2018). Thus, integrating perturbation phenotypes and cellular trajectories may inform the overall phenotypic impact of precise molecular changes. More generally, these data could provide new insights into the variability of phenotypes that result from a specific perturbation.

In this study, we analyzed 63 genotypes using ∼4,300 single cells (∼70 cells/genotype) in order to over-shoot the number of single-cells and sequencing depth required to reliably measure the effects of perturbations on the chromatin state. We suggest that future Perturb-ATAC screens target a lower number of cells per genotype (20), in order to adequately measure epigenomes, while expanding the number of genotypes assayed in a single experiment. Using our microfluidic platform that enables analysis of ∼100 cells per chip, this experimental strategy should easily enable screening of 150 genotypes in 3,000 cells. However, we do foresee several technological improvements that will enable higher throughput and more complex experimental designs. For example, this methodology should be easily adaptable for emerging methods of high-throughput scATAC-seq, such as using single-cell FACS sorting (Chen et al., 2018b, 2018a), the split-pool approach (Cusanovich et al., 2015, 2018), or nano-well or droplet-based methods (Mezger et al., 2018). These adaptations may be particularly useful in heterogeneous tissue contexts where cells occupy a larger diversity of chromatin states.

We designed the Perturb-ATAC platform to be compatible with widely-used CRISPR constructs, and therefore, we hope that it will be easily adoptable to existing screens in diverse systems. The ability to detect targeted RNA sequences could be easily adapted to detect alternative CRISPR constructs and classes of targets, such as orthogonal CRISPR guide RNAs, which could simultaneously regulate target gene activation and repression in the same cell (Boettcher et al., 2018; Najm et al., 2018). In addition to perturbation of *trans*-regulatory factors (TFs, chromatin regulators, and non-coding RNAs), this system could also be used to target individual *cis*-regulatory elements, for example to study the local effects of individual enhancer activation or repression on the expression of a given gene. This approach may be useful to finely dissect particular loci where both *cis*-regulatory elements and noncoding RNA transcripts have been shown to have functional effects on gene expression (Engreitz et al., 2016; Cho et al., 2018). Altogether, Perturb-ATAC provides a high-throughput platform to link genotypes with epigenetic phenotypes at single-cell resolution and ultimately reveal new molecular mechanisms that govern cell fate and function through modulation of the chromatin state.

## Acknowledgments

We thank members of the Khavari, Chang, and Greenleaf laboratories for helpful discussions. This work was supported by the US Veterans Affairs Office of Research and Development (P.A.K.), the National Institutes of Health (NIH) AR45192 (P.A.K.), P50-HG007735 (H.Y.C. and W.J.G.), R35-CA209919 (H.Y.C.), Parker Institute for Cancer Immunotherapy (A.T.S. and H.Y.C.), and Scleroderma Research Foundation (H.Y.C.). A.J.R was supported by a Stanford Bio-X Fellowship. K.R.P was supported by a Stanford Graduate Fellowship. A.T.S. was supported by a Parker Bridge Scholar Award from the Parker Institute for Cancer Immunotherapy and a Career Award for Medical Scientists from the Burroughs Wellcome Fund. W.J.G is a Chan Zuckerberg Biohub investigator. H.Y.C. is an investigator of the Howard Hughes Medical Institute.

## Author Contributions

A.J.R., A.T.S., H.Y.C., and P.A.K. conceived the project. A.J.R., K.R.P., A.T.S, Y.Q., B.W., A.J.O, D.S.K., M.R.M, and A.L.J. performed experiments and analyzed data. P.A.K., H.Y.C, and W.J.G. guided experiments and data analysis. A.J.R., K.R.P., A.T.S, H.Y.C., and P.A.K. wrote the manuscript with input from all authors.

## Supplementary Figure Legends

**Supplemental Figure 1: Perturb-ATAC CRISPRi construct and guide barcode detection scheme. (a)** Schematic of lentiviral plasmid encoding sgRNAs for CRISPRi as well as selection marker containing guide barcode. Stepwise targeted reverse transcription and PCR steps are displayed from top to bottom. **(b)** Overview of computational pipeline taking sequencing reads for GBC and producing final table of guide calls for each cell. **(c)** Detail on derivation of filtering parameters for per-cell sequencing depth and background reads. Left: distribution of reads aligning to any guide barcode are displayed for each of three representative plates. Middle: distribution of reads after plate-specific depth adjustment for high mode, resulting in uniform median depth for high mode across plates and uniform filter threshold of 1,000 normalized reads per cell. Right: Distribution of reads per cell not assigned to two most abundant guides, for cells annotated as single cell or doublet capture. Doublet wells separate into two modes, allowing determination of threshold separating unexpected high background in single capture wells.

**Supplemental Figure 2: Analysis of CRISPR sgRNA performance and consistency of Perturb-ATAC data cross separate C1 chips. (a)** Bar plots indicating the count of sgRNA sequence mismatch for random guides or guides selected for Perturb-ATAC. **(b)** Left: Description of workflow to calculate predicted off-target CRISPRi activity based on contribution of mismatches. Right: Histogram of predicted relative off-target activity for all sgRNAs used in this study, including up to 4 mismatches. **(c)** qPCR validation of CRISPRi gene expression knockdown after transduction with sgRNAs targeting the specified gene. **(d)** Bar plots indicating categories of sgRNA mismatch loci based on ATAC peak proximity and observed accessibility compared to non-targeting cells. **(e)** tSNE plots of all cells assayed in GM12878 experiment based on chromVAR feature deviation z-scores. For each plot, the cells assayed on a particular plate are highlighted.

**Supplemental Figure 3: Identification of differentially accessible genomic features in GM12878 screen. (a)** Violin plots of single cell accessibility relative to mean accessibility in non-targeting cells for significantly altered features in either EBER1, EBF1, EZH2, or SPI1 targeted cells. Each point represents an individual genomic feature (collection of genomic regions sharing an annotation such as a TF motif or ChIP-seq peak) in an individual cell. A maximum of 50 features are shown per genotype. **(b)** Scatter plots of accessibility in knockdown conditions, NFKB1 versus RELA (left) or EBER1 versus EBER1 (right). **(c)** Volcano plots for each single perturbation condition comparing perturbed cells to non-targeting control cells. Each point represents a genomic feature; significance threshold of FDR <= 0.025.

**Supplemental Figure 4: Inferred nucleosome profiles at differentially accessible regions. (a)** Schematic depicting generation of short (<100bp) ATAC fragments from sub-nucleosome regions and large fragments (180-247bp) spanning nucleosome-protected regions. **(b)** Metaplots of sub-nucleosome and nucleosome fragment signal at CTCF motif regions overlapping with CTCF ChIP-seq peaks in GM12878. Signal represents average of two non-targeting cell populations, gray range represents standard deviation between samples. **(c)** Metaplots of sub-nucleosome and nucleosome signal at differentially accessible regions.

**Supplemental Figure 5: Expanded analysis of perturbed intercellular genomic feature correlation networks. (a)** Heatmap of correlation matrices for genomic features. Values indicate Pearson correlation across non-targeting cells for accessibility of two genomic features. Ward’s hierarchical clustering was used to identify five modules with substantial intra-cluster correlation. **(b)** Listing of key features in each module. **(c)** Heatmap of correlation matrix for genomic features in IRF8 knockdown cells. **(d)** Left: Box plots of single cell accessibility for CTCF and SMAD5 features in non-targeting and DNMT3A knockdown cells. Right: Histogram of z-score of number of altered correlations for each feature in DNMT3A knockdown cells. **(e)** Heatmap of difference in feature correlations between NFKB1 knockdown cells (bottom) and RELA knockdown cells (top). **(e)** Heatmaps of feature correlations for Module 1 vs. Module 5 in non-targeting cells or EBER2 knockdown cells. **(f)** Histogram of change in feature correlations for SPI1 knockdown versus non-targeting 1 cells, used to inform thresholds for designation of altered correlation. **(g)** Table of counts and highlighted top altered-correlation features based on 5% FDR threshold.

**Supplemental Figure 6: Perturb-ATAC CRISPR KO constructs and activity. (a)** Schematic of lentiviral plasmids for sgRNA and Cas9 expression. **(b)** Sanger sequencing traces of the 100bp surrounding sgRNA 3’ end for each target gene. Sequencing proceeded in forward direction (left to right), resulting in abrupt drop in sequencing alignment after sgRNA due to mixture of indels.

**Supplemental Figure 7: Perturb-ATAC CRISPR KO direct guide detection scheme. (a)** Schematic of lentiviral plasmid encoding sgRNA for CRISPR knockout. Stepwise targeted reverse transcription and PCR steps are displayed from top to bottom. **(b)** Distributions of reads per cell mapping to a sgRNA variable sequence. For each plate, a clear high mode of reads was identified and used to determine a depth cutoff. **(c)** Distribution of proportion of all reads per cell mapping to known sgRNA sequence. **(d)** Distribution of proportion of reads per cell associated with background (third most common) guide sequence. Cells in low mode passed filter. **(e)** For cells passing previous filters, distribution of proportion of reads associated with second most common guide. Cells in the low mode of this distribution were considered to express a single guide, while cells in the high mode were considered to express two guides. **(f)** Scatter plots of proportion of reads associated with two guide sequences for all cells passing final filters.

**Supplemental Figure 8: Altered features in keratinocyte differentiation induced by genetic perturbations. (a)** Signal track indicating a ZNF750 binding site that gains accessibility in targeted cells, indicating repressive activity of ZNF750. **(b)** Scatter plot of principal component (PC) values for unperturbed keratinocytes. PC space was generated using altered features from specific single TF knockout cells. Yellow line represents pseudotime trajectory connecting centroids of cells from each differentiation day. (**c**) Scatter plot of PC values for all perturbed and non-targeting cells embedded in PC space generated in (a). Cells are scored and colored by progression along pseudotime trajectory. These pseudotime values were used to assess the enrichment or depletion of knockout versus non-targeting cells in Figure 7F. (**d**) As in Figure 8B, scatter plots of observed versus expected (based on additive model) accessibility in double knockout cells. **(e)** Scatter plot of absolute log2 fold changes of features in single knockout cells versus double knockouts (r ∼ 0.18).

## CONTACT FOR REAGENT AND RESOURCE SHARING

Further information and requests for resources and reagents should be directed to and will be fulfilled by the Lead Contact, Paul Khavari.

## METHOD DETAILS

### CRISPRi targeting in GM12878

To generate the Perturb-ATAC vector with guide barcodes used in the GM12878 experiments, we modified previously-described CRISPRi vectors (Adamson et al., 2016; Cho et al., 2018). Briefly, we designed three sgRNAs per target gene, each targeting a different region between the transcriptional start site and 200 nucleotides into the gene body. One sgRNA each was cloned into pMJ114 (bovine U6, Addgene, Cat#85995), pMJ117 (human U6, Addgene, Cat#85997) or pMJ179 (mouse U6, Addgene, Cat#85996), digested with BstXI and BlpI, using NEBuilder Hifi DNA Assembly Master Mix. Then the respective U6 promoter and sgRNA sequences were amplified by PCR and assembled into the lentiviral vector (digested using XbaI and XhoI) using NEBuilder Hifi DNA Assembly Master Mix. Subsequently, individual colonies for each 3x sgRNA plasmid were digested using PciI and EcoRI, and a randomized 22 bp barcode (ordered from IDT as 5’- [overhang][NNN…][overhang]-3’) was assembled with NEBuilder Hifi DNA Assembly Master Mix. The sgRNA sequences and GBC sequences of all plasmids were confirmed by Sanger sequencing.

To generate CRISPRi virus, we used HEK 293T cells maintained in DMEM with 10% FBS, 1% Pen-Strep. Cells were seeded at 4 million per 10cm dish, and the following day transfected with 4.5ug pMP.G, 1.5ug psPAX2, and 6ug sgRNA vector using OptiMEM and Lipofectamine 3000. Two days later, the supernatant was collected and filtered with a 0.44 μm filter, and virus was concentrated 1:10 using Lenti-X Concentrator (Clontech).

GM12878 maintained in RPMI 1640 (Thermo Fisher) with 10% FBS and 1% Penicillin-Streptomycin (Thermo Fisher) were then seeded at 300,000 cells per well of a 6-well plate and 40ul of concentrated virus was added to the media the following day. Two days later, we exchanged the media for media containing 1ug/ml puromycin to select for the sgRNA vector. Selection media was refreshed on day five, and on day seven cells selection media was exchanged for regular media (containing no puromycin) and cells were either assayed or frozen in viable conditions with BamBanker cryopreservation media. Cells were sorted by flow cytometry for viability and expression of mCherry before being assayed by Perturb ATAC-seq. Cells were maintained between 200,000 and 1 million per mL. RNA was extracted with Trizol and purified using Qiagen RNeasy columns, and gene expression knockdown was confirmed using the Agilent Brilliant II qRT-PCR 1-Step kit. qRT-PCR was performed in duplicate, and expression values for each sample were normalized against 18S. Gene expression values for CRISPRi are reported as average fold change against both non-targeting control samples.

### Culture, differentiation, and CRISPR knockout in primary keratinocytes

Primary human keratinocytes were isolated from fresh, surgically discarded neonatal foreskin. Keratinocytes were grown in 1:1 KCSFM and Medium 154 (Life Technologies). Keratinocytes were induced to differentiate by addition of 1.2mM calcium for 3 or 6 days in full confluence.

We generated custom Cas9 and sgRNA expression vectors for CRISPR knockout in keratinocytes. For Cas9 expression, we amplified the Cas9 gene from the lentiCRISPRv2 vector (Sanjana et al., 2014) and cloned this fragment into pLex-MCS (Thermo Fisher) along with a fusion P2A-blasticidin resistance cassette in exchange for the IRES-puromycin resistance cassette in pLex-MCS. For sgRNA expression, we modified the sgRNA F+E scaffold (Chen et al., 2013; https://www.addgene.org/59986/) in two ways. First, we exchanged the murine U6 promoter and telomerase-targeting sgRNA with the human U6 promoter, stuffer region, and associated BsmBI cloning sites from lentiCRISPRv2. Additionally, we removed a BsmBI restriction site in the puromycin resistance gene by introducing a non-synonymous mutation.

To generate lentivirus, we seeded 400,000 HEK 293T cells into a single well of a 6-well dish, and the following day we transfected either our Cas9 vector or sgRNA vector (1.3 ug) along with pMDG (0.3 ug) and p8.91 (1 ug) using Lipofectamine 3000 (Thermo Fisher). Supernatant was collected at 48hrs and 72 hrs, filtered through a 0.45um PES membrane, and concentrated to a pellet with Lenti-X Concentrator. One unit of Cas9 virus corresponded to the concentrated supernatant from one 6-well of HEK 293T. One unit of sgRNA virus corresponded to one eighth of the concentrated supernatant from one 6-well of HEK 293T.

Primary keratinocytes were seeded at 300,000 cells per well of a 6-well dish along with one unit of Cas9 virus and polybrene (0.1 ug/ml). After one day, two wells were harvested, mixed, and expanded into a 15cm dish containing normal culture media with 2ug/ml blasticidin. After four to six days of selection, cells were again seeded at 300,000 cells per well of a 6-well dish along with one unit of sgRNA virus and polybrene (0.1 ug/ml). After one day, one well was harvested and transferred to a 15cm dish containing normal culture media, puromycin (1 ug/ml) and blasticidin (2 ug/ml). After six days of selection, cells were seeded at highVconfluence with 1.2 mM calcium for differentiation. Cells were harvested after three days of differentiation and viably frozen in culture media with 10% DMSO.

Cas9 nuclease activity was assessed by PCR amplifying ∼800bp fragments of cDNA surrounding sgRNA binding sites and analyzing the resulting fragments by Sanger sequencing (oligo sequences in Supplementary Table 6). Images depicted in Figure S6 were generated using Geneious 7.1.4. cDNA was generated by extracting RNA from cells with the RNeasy Mini Kit (Qiagen) and performing reverse transcription with the iScript cDNA Synthesis Kit (Bio-Rad).

### Bulk ATAC-seq

Cells were isolated and subjected to ATAC-seq as previously described (Corces et al., 2017). Briefly, 50,000 cells were pelleted after sorting and resuspended in 50ul of ATAC resuspension buffer (RSB) with 0.1% NP40, 0.1% Tween-20, and 0.01%. After three minutes, 1ml of ATAC RSB with 0.1% Tween-20 was added, tubes were inverted, and nuclei were centrifuged at 500 rcf for 10 min. Supernatant was carefully removed and nuclei were resuspended in 50ul transposition mix (25ul TD buffer, 2.5ul transposase, 16.5ul PBS, 0.5ul 0.1% digitonin, 0.5ul 10% Tween-20, and 5ul water). Transposition was performed for 30 minutes at 37 C with shaking in a thermomixer at 1000 RPM. Reactions were purified with a Zymo DNA Clean & Concentrator 5 kit and library generation was performed as described previously (Corces et al., 2017).

### Single-cell ATAC-seq

Single-cell ATAC-seq was performed as previously described (Buenrostro et al., 2015). In brief, cells were sorted by flow cytometry for viability and to remove cell aggregates. The C1 Single-Cell Auto Prep System was used with the Open App^™^ program (Fluidigm, Inc.). The Open App scripts from the “ATAC Seq” collection from Fluidigm were used to prime the C1 IFC microfluidic chip, load cells, and run the ATAC sample prep protocol. Fluidigm scripts are available from Fluidigm Script Hub, https://www.fluidigm.com/c1openapp/scripthub.

### Perturb ATAC-seq

#### Cell isolation and microfluidic reactions on the IFC

We adapted the C1 Single-Cell Auto Prep System with its Open App^™^ program (Fluidigm, Inc.) to perform Perturb-ATAC-seq. C1 IFC microfluidic chips were first primed by following the Open App script “Biomodal Single-Cell Genomics: Prime”. Single cells were then captured using the Fluidigm Open App script “Biomodal Single-Cell Genomics: Cell Load.” GM12878 or keratinocyte cells were first isolated by FACS sorting and then washed three times in C1 DNA Seq Cell Wash Buffer (Fluidigm). Cells were resuspended in DNA Seq Cell Wash Buffer at a concentration of 300 cells/μL and mixed with C1 Cell Suspension Reagent at a ratio of 3:2 (cells:reagent). 15 μl of this cell mix was loaded onto the IFC. After cell loading, all wells were visualized by imaging on a Leica CTR 6000 microscope to identify captured cells.

Cells were then subjected sequentially to lysis and transposition, transposase release, quenching with MgCl_2_, reverse transcription, and PCR, using the custom Open App IFC script “Biomodal Single-Cell Omics: Sample Prep.” For lysis and transposition, 30μL of Tn5 transposition mix was prepared (22.5μL 2x TD buffer, 2.25μL transposase (Nextera DNA Sample Prep Kit, Illumina), 2.25μL C1 Loading Reagent without salt (Fluidigm), 0.45μL 10% NP40, 2.25μL SuperaseIN RNase inhibitor, and 0.3μL water). For transposase release, 20μL of Tn5 release buffer mix was prepared (2μL 500 mM EDTA, 1μL C1 Loading Reagent without salt, and 17μL 10 mM Tris-HCl Buffer, pH 8). For MgCl_2_ quenching, 20μL of MgCl_2_ quenching buffer mix was prepared (18 μL 50 mM MgCl2, 1μL C1 Loading Reagent without salt, and 1μL 10 mM Tris-HCl Buffer, pH 8). For reverse transcription, 30μL of RT mix was prepared (15.55μL H_2_0, 3.7μL 10x Sensiscript RT buffer (Qiagen), 3.7μL 5 mM dNTPs, 1.5μL C1 Loading Reagent without salt (Fluidigm), 1.85μL Sensiscript RT (Qiagen), and 3.7μL 6 μM RT primer mix (6uM each of V1 GBC sequencing oligos or 6uM each of V1 sgRNA sequencing oligos, see Supplementary Tables 3 and 6 for oligo sequences). Finally, for ATAC and GBC/sgRNA PCR, 30uL of PCR mix was prepared (8.62μL H_2_0, 13.4μL 5x Q5 polymerase buffer (NEB), 1.2μL 5 mM dNTPs, 1.5μL C1 Loading Reagent without salt, 0.67μL Q5 polymerase (2U/μL; NEB), 0.8μL 25 μM non-indexed custom Nextera ATAC-seq PCR primer 1, 0.8μL 25 μM non-indexed custom Nextera ATAC-seq primer 2, and 3 μL 6 μM GBC or sgRNA primer mix.

7 μL lysis and transposition mix, 7 μL transposase release buffer, 7 μL MgCl_2_ quenching buffer, 24 μL RT mix, and 24 μL PCR mix were added to the IFC inlets. On the IFC, Tn5 lysis and transposition reaction was carried out for 30 minutes at 37^°^. Next, transposase release was carried out for 30 min at 50^°^C. MgCl_2_ quenching buffer was immediately added and chamber contents were immediately incubated with RT mix for 30 minutes at 50^°^C. Finally, gap filling and 8 cycles of PCR were performed using the following conditions: 72^°^C for 5 min and then thermocycling at 94^°^C for 30s, 62^°^C for 60s, and 72^°^C for 60s. The amplified transposed DNA was harvested in a total of 13.5 μL C1 Harvest Reagent. Following completion of the on-chip protocol (∼4-5hrs), chamber contents were transferred to 96-well PCR plates, mixed, and divided for further amplification of ATAC-seq fragments (6-7 *μ*l) or GBC/sgRNA fragments (6.5 *μ*l).

For method development and RT primer troubleshooting, the Perturb-ATAC-seq protocol can be exactly scaled 1000x and performed on 1000 cells in Eppendorf tubes. Following lysis, transposition, and transposase release, RNA can be reverse-transcribed and subjected to PCR amplification to check the amplification efficiency and specificity of a chosen primer set.

#### Amplification of ATAC-seq libraries

∼7 μL of harvested libraries were amplified in 50 μL PCR for an additional 15 cycles with 1.25 μM Nextera dual-index PCR primers in 1x NEBnext High-Fidelity PCR Master Mix using the following PCR conditions: 72^°^C for 5 min; 98^°^C for 30s; and thermocycling at 98^°^C for 10s, 72^°^C for 30s, and 72^°^C for 1 min. The PCR products were pooled and purified on a single MinElute PCR purification column (Qiagen). Libraries were quantified using qPCR (Kapa Library Quantification Kit for Illumina, Roche) prior to sequencing using 2x76bp paired-end reads on an Illumina NextSeq 550 or 2x75bp reads on an Illumina MiSeq.

#### Amplification of guide barcode and guide RNA sequencing libraries

Three rounds of off-C1 PCR were performed to generate GBC and sgRNA sequencing libraries (See Supplementary Tables 3 and 6 for V1,V2, and V3 oligo sequences). First (V1 PCR), 6.5ul of harvested libraries were amplified in a 20 ul PCR (harvested DNA with 10ul NEBNext Master Mix, 0.1 ul of each V1 primer at 200uM, and remaining volume of water). Reactions amplified for 17 cycles with the following parameters: 98 C for 30s, then cycling of 98 C for 10s, 63 C for 30s, and 72 C for 45s, followed by 72 C for 5 min. Second, 2ul of the V1 PCR product (without purification) was transferred to a subsequent 20ul reaction with 10ul NEBNext Master Mix, 0.1 ul of each V2 primer at 200uM, and remaining volume of water. Reactions were amplified for 15 cycles using the same parameters used for V1 reactions. A final 20ul V3 cell indexing PCR was performed using 2ul of the V2 reaction product, 2ul each of Illumina Indexing primers at 10 uM, 10ul NEBNext Master Mix, and the remaining volume of water. Reactions were amplified for 15 cycles using the same parameters used for V1 and V2 reactions.

Finally, V3 reactions were pooled and purified using the Qiagen MinElute kit. Libraries were further purified by size selection on polyacrylamide gel electrophoresis (6% TBE Novex gel, Thermo Fisher). Libraries were mixed with BlueJuice loading dye (Thermo Fisher), run for 35 min at 160 V and visualized using SybrSafe stain (Thermo Fisher), using 5ul of stain in 30ml of TBE running buffer for 10 min. Gels were visualized on a blue-light transilluminator and slices in size range for GBC library fragments (289 bp) or sgRNA library fragments (232 bp) were cut using a scalpel. Gel slices were placed in a 0.75ml tube with a hole punctured in the bottom using a syringe, and this tube was placed in a 1.5ml DNA LoBind tube (Eppendorf). These tubes were centrifuged for 3 min at 13k RPM to crush the gel slice, then 300ul Salt Crush Buffer (500 mM NaCl, 1 mM EDTA, 0.05% SDS) was added and this mix was incubated at 55 C overnight in a thermomixer with 1000 RPM shaking. The next day, samples were cooled to RT, centrifuged through a Spin-X column (one minute, 13k RPM), and purified with a Zymo DNA Clean & Concentrator 5 kit. Libraries were quantified by qPCR (Kapa Library Quantification Kit for Illumina, Roche) before sequencing on an Illumina MiSeq at 10-14 pM final concentration with 15-40% PhiX.

## QUANTIFICATION AND STATISTICAL ANALYSIS

### Single cell and bulk ATAC primary processing and chromVAR analysis

Single cell and bulk ATAC read alignment, quality filtering, and duplicate removal were performed was previously described (Buenrostro et al., 2015). Briefly, adapter sequences were trimmed, sequences were mapped to the hg19 reference genome using Bowtie2 (Langmead and Salzberg, 2012; and the parameter -X2000), and PCR duplicates were removed using Picard Tools. Reads mapping to the mitochondria were discarded for further analysis. We observed an extremely low rate of ATAC reads matching the CRISPR viral construct (median 0.0049%) and found no evidence of the abundance of CRISPR construct matching reads influencing epigenomic profiles.

Single cell ATAC-seq calculation of TF deviation was performed using chromVAR (in R, version 1.1.1; Schep et al., 2017). Briefly, for each TF, ‘raw accessibility deviations’ were computed by subtracting the expected number of ATAC-seq reads in peaks for a given motif from the observed number of ATAC-seq reads in peaks for each single cell. Expected reads were calculated from the population average of all cells for the GM12878 experiment and unperturbed cells only for the keratinocyte experiment. This value is subtracted by the mean deviation calculated for sets of ATAC-seq peaks with similar accessibility and GC content to obtain a bias-corrected deviation value, and additionally divided by standard deviation of the deviation calculated for the background sets to obtain a Z-score.

For the GM12878 experiments, we used a set of peaks derived from DNAse I hypersensitivity data (downloaded from http://genome.ucsc.edu/cgi-bin/hgTrackUi?db=hg19&g=wgEncodeUwDnase) from a broad variety of hemopoietic cell lines (all GM lines, HL-60, Th1, Jurkat, K562) plus additional lines (HepG2, HUVEC, NHEK), to account for the possibility of opening peaks outside the blood lineage. These peaks were each filtered against the wgEncodeDacMapabilityConsensusExcludable.bed blacklist, sorted by intensity, and the top 75,000 peaks for each sample were merged. These peaks were then centered and resized to 1kb uniform peaks (238,349 final peaks).

For the keratinocyte experiment, we merged peaks called on bulk ATAC-seq from undifferentiated cells and cells differentiated for three or six days. Peaks were called using the MACS2 command *macs2 callpeak --nomodel –nolambda –-call-summits --shift -75 --extsize 150* (Zhang et al., 2008). First, peaks with q-value < 0.01 from each day were merged. In the case of overlapping peaks, the summit associated with the lowest q-value was selected as the merged peak summit, and the 1kb window centered on that summit was used as the uniform peak for chromVAR (94,633 final peaks).

For GM1878 analysis, narrowPeak ChIP-seq files (optimal IDR thresholded peaks) were downloaded from ENCODE and imported as supplementary annotations in chromVAR. Prior to use, these files were filtered against the wgEncodeDacMapabilityConsensusExcludable.bed blacklist. H3K27me3 and H3K27ac narrowPeak files for different tissues were downloaded from the Roadmap Epigenomics website (http://www.roadmapepigenomics.org/data/).

### Guide barcode sequencing analysis for GM12878 experiments

For GM12878 experiments, raw reads for GBC libraries were matched to a list of GBC sequences to generate a table of counts for each cell and each GBC analyzed in the experiment (see Figure S1, custom scripts written in Python available upon request). First, any read not containing the expected 27 nt sequence prior to the GBC was discarded, allowing for a maximum Levenshtein distance of 2 to account for sequencing errors. The subsequent 22 nt sequence was then compared to a list of GBC sequences, allowing for a maximum Levenshtein distance of 3 to be considered a match. Note that the minimum Levenshtein distance between any two of our GBC sequences was 10. This generated a counts-per-cell table for each GBC sequence and cell.

This table was normalized for read depth by plate by assessing the maximum density of log-transformed counts using the scipy.stats.gaussian_kde function (see Figure S1C). This distribution exhibits a bimodal distribution corresponding to wells with productive and unproductive GBC detection. A normalized GBC read cutoff of 1000 reads/cell was set (Figure 2A, this was empirically determined based off the separation between wells with and without a cell capture). Cells displaying high background reads, as determined by having greater than 0.005 proportion reads not aligning to the top two GBC sequences, were further filtered (this cutoff was set from empirical observations of “background” in doublet wells, which are expected to contain up to four GBC sequences; Figure S1C). We distinguished cells expressing a single or double sgRNAs based off the percent of reads aligning to the second-most common GBC (single, <1% double, >5%). This workflow resulted in far more double-targeted cells than would be observed solely from the observed doublet rate calculated from the appearance of double GBC-expressing cells in our initial single-targeting experiment (∼ 2.9%). tSNE plots shown in Figure S2 were generated using the manifold.TSNE function in the Python package scikit-learn.

We empirically determined a target minimum cell number required for analysis by down-sampling cells from a larger pool and comparing accessibility profiles. This analysis indicated that the vast majority of samples of five cells were highly correlated (r > 0.8) with a bulk ATAC-seq profile. Additionally, previous reports have shown that aggregation of five or more cells is sufficient to accurately reproduce chromatin accessibility profiles (Satpathy et al., 2018; Schep et al., 2017). In line with these findings, we designed Perturb-ATAC experiments to yield the maximal number of genotypes supported by at least five cells; indeed 38/40 genotypes for GM12878 cells and 23/23 genotypes for keratinocytes consist of greater than five cells.

### Direct sgRNA sequencing and analysis for keratinocyte experiments

For keratinocyte experiments, raw reads for sgRNA sequencing were matched to a list of sgRNA sequences used in the experiment. We required strict matching of the 20bp variable sequence along with 18bp of the standard sgRNA backbone. Matching was performed with custom scripts (available upon request) and resulted in the counts-per-cell table for each sgRNA.

We then normalized this table for read depth by assessing the plate-specific distribution of log-transformed total counts per cell (**Figure S7**). The collection of counts per cell exhibited a bimodal distribution likely corresponding to productive and failed sgRNA detection. We drew a cutoff in between the two modes as a first filter, and further required cells to exhibit low background (reads associated with the third most common sgRNA in each cell). Cells with greater than 1% of reads associated with background were excluded from analysis. Finally, we distinguished cells expressing one or two sgRNAs based on the distribution of proportions of reads associated with the second most common sgRNA in each cell. Cells with fewer than 1% of reads associated with the second most common sgRNA formed a clear mode in this distribution and were considered to express only the most common sgRNA, while cells with greater than 10% of reads associated with the second most common sgRNA were considered to express both the first and second most common sgRNAs.

### Identification of differentially accessible genomic features and regions

We generated an empirical null distribution of accessibility values for each feature in order to assess the significance of any observed difference between mean accessibility in a set of perturbed cells compared to cells expressing non-targeting control sgRNAs. For each genomic feature (peak or chromVAR motif/annotation), we first calculated the median deviation z-score (for chromVAR features) or fragment counts (for peaks) in cells expressing each sgRNA or combination of sgRNAs. Cells expressing a targeting sgRNA in combination with a non-targeting sgRNA were analyzed with targeting sgRNA-only cells. With the goal of assessing the null hypothesis that targeting and non-targeting cells exhibit the same accessibility, we pooled equal numbers of cells from targeting and non-targeting cells. This population was then randomly divided into two sets by permuting the cell-genotype labels, and the permuted median accessibility difference of these two populations were compared to the observed median accessibility difference. This process was repeated 5000 times to generate a null distribution, and the rate of detecting a median accessibility difference as extreme or greater in the null distribution compared to the observed targeting cells was reported as the false discovery rate (FDR).

Differentially accessible regions were found using a similar approach with the exception that we limited the set of total regions under consideration to those exhibiting at least one read per five cells in one of the conditions under consideration for each comparison. Genome browser tracks of differentially accessible regions were generated by pooling cells associated with a particular sgRNA genotype. We first generated bedGraph files scaled to 500,000 reads using the genomeCoverageBed tool (BedTools v2.17.0) then generated bigWig files using the bedGraphToBigWig tool from UCSC (http://hgdownload.soe.ucsc.edu/admin/exe/). Tracks were finally displayed in the WashU Epigenome Browser.

### Statistical analysis of SPI1 motif-containing region accessibility in SPI1-depleted cells

For Figure 2G, we determined an empirical false discovery rate for the observed changes in SPI1 motif region accessibility. For bulk-ATAC and Perturb-ATAC samples separately, we calculated the z-score of the SPI1 motif accessibility change in perturbed cells compared to all other features. Then to generate a null distribution, we permuted the sample labels between Non-targeting #1, Non-targeting #2, and SPI1-targeting 1000 times and in each trial recorded the z-score of SPI1 motif change in accessibility compared to the non-targeting controls. In this analysis, for both bulk-ATAC and Perturb-ATAC, no trial yielded a result as extreme as the result observed in the unpermuted sample.

### Inferred nucleosome and sub-nucleosome profiles and score calculation

The aggregate profiles of nucleosomal signals at differentially accessible regions were derived from total ATAC fragments as described previously (Bao et al., 2015). Briefly, ATAC fragments sized 180-247bp were considered nucleosome-spanning and used to infer positions of nucleosomes in aggregate locus profiles (metaplots). Differentially accessible regions were centered based on the signal summit as identified by Macs2 (using the flags –-call-summits --shift-75 --extsize 150) and filtered for an FDR < 0.1 and log2 fold change > 1. We then calculated the fragment count in 10bp windows spanning 1000 bp upstream and downstream of the region summit. These profiles were normalized to the average signal in the 25 downstream windows to account for sequencing depth and the resulting enrichment values were smoothed in R using the smooth.spline() function with parameter spar = 0.5.

To quantify the presence of peak central versus flanking nucleosome in each metaplot, we calculated the ratio of flanking nucleosome signal density (-180 to -80bp relative to peak summit and +80 to +180bp relative to peak summit) to central nucleosome signal density (-20 to +20bp relative to peak summit). We report this ratio as the central nucleosome score.

### Analysis of inferred regulatory networks

To identify sets of genomic features whose activities were correlated across single cells, suggestive of shared regulatory relationships, we computed the Pearson correlation of each feature with each other feature across all single cells of a given genotype. Only features that were significantly altered in at least one genotype were considered, and redundant annotations were removed, resulting in 390 motif/ChIP feature annotations for analysis. Ward’s hierarchical clustering was performed and features displaying low intra-cluster correlation were excluded from further analysis (Figure S5A). The modules shown in subsequent analysis were defined based off Ward’s hierarchical clustering of the remaining features in non-targeting cells. Clustering was performed using the Seaborn clustermap function using Ward’s method for clustering.

For each Perturb-ATAC genotype, the feature-feature correlation across single cells was computed. The difference in correlation between a given genotype and non-targeting cells was computed by subtracting the Pearson correlation in the respective genotype from non-targeting cells. A permutation test was used to assess the significance of the observed change in correlation for any pair of features. For each genotype, the same number of cells was randomly sampled from all perturbed cells 10,000 times, and the changes in correlation in the randomly sampled cells relative to non-targeting cells were used to create a null distribution for each feature-feature pair (in each genotype). A 5% cutoff was used to call significantly altered correlations. To quantify module-level changes in regulatory relationships, we quantified the percent of all feature-feature pairs in a given module whose correlations were significantly altered.

### Analysis of epistasis for accessibility of genomic features

We assessed the degree of epistasis in double perturbation conditions by comparing observed phenotypes in double perturbation conditions to phenotypes expected based on a model of non-interaction. For this analysis, we scored the accessibility of genomic features based on the sum of raw reads accumulating in peaks associated with that feature in each cell. Feature counts were normalized by the total number of reads for features in each cell and log2-transformed with the addition of a pseudocount. For each collection of cells sharing a genotype, the mean value of log2 counts was compared to the mean value of log2 counts for a mix of cells expressing non-targeting sgRNAs, resulting in a log2 (fold change of perturbation vs. non-targeting). The additive expectation was based on a multiplicative model of non-interaction, (i.e., CRISPR AB = CRISPR A x CRISPR B), which we calculated by adding the single perturbation fold changes in log2-space. For each genomic feature, the degree of interaction (difference between observed accessibility change and that expected under the non-interaction model) was calculated.

To identify generally additive vs. non-additive features (Figure 5D,E), the interaction degree was averaged across perturbations. To compute the permuted background, we permuted the single-double pairings by randomly choosing a double sgRNA genotype and two random single sgRNA genotypes. The difference between the “expected” change (based on the two random sgRNA genotypes) and the “observed” changed (based on the random double sgRNA genotype) was then computed. This process was repeated once for each double sgRNA genotype observed in our dataset.

We further categorized features as additive, synergizing, and buffering for a particular interaction (Figure 8) by comparing the observed degree of interaction to a null distribution generated by permuting cell identities. This procedure was performed separately for each feature to account for differences in scale and variability across features. The null distribution was generated by randomly sampling three pools of cells from all perturbed cells: a null double perturbation set, and two null single perturbation sets. The difference between observed double perturbation phenotype and the expected value from the non-interaction model was calculated, and this procedure was repeated 1000 times. Genotypes exhibiting interaction degrees beyond 95% of the null values were considered interacting. Interactions in which the double phenotype had a more extreme magnitude than expected were labeled synergistic, while others were labeled buffering.

### Analysis of tissue H3K27me3 and autoimmune-associated SNPs

128 consolidated narrowPeak files for H3K27me3 peaks (corresponding to different tissues/cell-types) were downloaded from the Roadmap Epigenomics Consortium website. Peaks that were found across at least 30 samples were considered common H3K27me3 peaks. Individual narrowPeak files were then filtered against this set of common H3K27me3 peaks, as well as the wgEncodeDacMapabilityConsensusExcludable blacklist. The resulting files were subsequently centered and resized to create uniform 1kb peaks, and imported into chromVAR as an annotation set. To identify peaks repressed in the GM12878 lineage but active in other tissues, H3K27ac narrowPeaks from blood tissues present in the Roadmap Epigenomics Consortium dataset were downloaded and intersected with the GM12878 H3K27me3 narrowPeak set using the bedtools intersect command. These were similarly filtered aginst the same blacklist, centered, and resized to create uniform 1kb peaks, and imported as a chromVAR annotation set.

SNPs associated with autoimmune diseases were downloaded from (Farh et al., 2015). These were aggregated by each autoimmune disease, and intersected with FitHiC calls (processed using 10kb genomic windows) from GM12878 H3K27ac HiChIP data (Mumbach et al., 2017). For each disease, the SNP (ultimately resized to a 10kb genomic window), as well as any windows in contact with that SNP, were aggregated to create a disease-specific chromVAR annotation set. As it is difficult to determine *a priori* whether a disease state would result from increased or decreased accessibility at a given site, we reported the absolute value change chromVAR deviation z-score for each genotype.

### Pseudotime calculation and identification of feature modules

For the keratinocyte experiment, the normal differentiation pseudotime trajectory was calculated using Monocle 2 (Qiu et al., 2017b). The feature deviation matrix including unperturbed and CRISPR knockout cells was first processed using Seruat 2.0.1 (Butler et al., 2018) to regress out plate and experiment batch effects. The Seurat function ScaleData was used (with parameters do.scale=F and do.center=F) to perform batch regression. To identify modules of dynamic features across differentiation, we first filtered for features that exhibited standard deviation greater than 1.3 in any comparison of normal differentiation conditions (Day 0, 3, or 6). Similar features associated with the AP1 motif were merged into a single feature. The matrix of these features vs. cells (arranged by increasing pseudotime) was hierarchically clustered using the heatmap.2 function in the gplots R package, resulting in three major clusters (referred to as modules).

Individual peaks approximately matching the kinetics of modules were identified in order to find associated genes (Figure 7C). Peaks exhibiting a log2 fold change less than 0.5 between conditions were considered stable and a fold change greater than 2 was considered dynamic. Peaks exhibiting decreased accessibility on both Day 3 and Day 6 (relative to Day 0) were considered Module 1 peaks. Peaks exhibiting increased accessibility on Day 3 versus Day 0 but stable accessibility between Day 6 and Day 0 were considered Module 2 peaks. Peaks exhibiting stable accessibility between Day 3 and Day 0 but gained accessibility on Day 6 versus Day 0 were considered Module 3 peaks. Genes were considered potential regulatory targets of a peak if the gene transcription start site fell within 50kb of the peak.

### Altered differentiation trajectory and module activity analyses

For each single perturbation in the keratinocyte experiment, a custom pseudotime was calculated in order assess the enrichment or depletion of cell occupancy along the differentiation trajectory (Figure 7F). ChromVAR deviations regressed for experimental batch effects and merged AP1 features were used for this analysis. Cells from each perturbation were pooled with non-targeting cells and a custom principal component analysis (PCA) space was generated. Features altered in each perturbation (FDR < 0.1, change in z-score > 0.25) were selected in order to achieve maximum separation of control and perturbed cells, and a PCA was generated with the R prcomp function (center=T, scale=T). Next, non-perturbed cells from all stages of differentiation were analyzed and a trajectory was calculated progressing from undifferentiated cells (Day 0) to mid-differentiation (Day 3) and finally late-differentiation (Day 6). The trajectory was determined by plotting a linear path between centroids of the three cell populations representing each stage of differentiation. Finally, the distribution of non-targeting cells and targeted cells was calculated along eight equally sized bins in this trajectory, and the log2 fold change of the proportion of cells in each been was reported as an enrichment.

## DATA AND SOFTWARE AVAILABILITY

Sequencing data (fastq files) and processed data files are available from GEO.

## References

Adamson, B., Norman, T.M., Jost, M., Cho, M.Y., Nuñez, J.K., Chen, Y., Villalta, J.E., Gilbert, L.A., Horlbeck, M.A., Hein, M.Y., et al. (2016). A Multiplexed Single-Cell CRISPR Screening Platform Enables Systematic Dissection of the Unfolded Protein Response. Cell 167, 1867-1882.e21.

Adamson, B., Norman, T.M., Jost, M., and Weissman, J.S. (2018). Approaches to maximize sgRNA-barcode coupling in Perturb-seq screens. bioRxiv 298349.

Arrand, J.R., Young, L.S., and Tugwood, J.D. (1989). Two families of sequences in the small RNA-encoding region of Epstein-Barr virus (EBV) correlate with EBV types A and B. J. Virol. 63, 983–986.

Bao, X., Rubin, A.J., Qu, K., Zhang, J., Giresi, P.G., Chang, H.Y., and Khavari, P.A. (2015). A novel ATAC-seq approach reveals lineage-specific reinforcement of the open chromatin landscape via cooperation between BAF and p63. Genome Biol. 16, 284.

Boettcher, M., Tian, R., Blau, J.A., Markegard, E., Wagner, R.T., Wu, D., Mo, X., Biton, A., Zaitlen, N., Fu, H., et al. (2018). Dual gene activation and knockout screen reveals directional dependencies in genetic networks. Nat. Biotechnol. 36, 170–178.

Boxer, L.D., Barajas, B., Tao, S., Zhang, J., and Khavari, P.A. (2014). ZNF750 interacts with KLF4 and RCOR1, KDM1A, and CTBP1/2 chromatin regulators to repress epidermal progenitor genes and induce differentiation genes. Genes Dev. 28, 2013–2026.

Buenrostro, J.D., Giresi, P.G., Zaba, L.C., Chang, H.Y., and Greenleaf, W.J. (2013). Transposition of native chromatin for fast and sensitive epigenomic profiling of open chromatin, DNA-binding proteins and nucleosome position. Nat. Methods 10, 1213–1218.

Buenrostro, J.D., Wu, B., Litzenburger, U.M., Ruff, D., Gonzales, M.L., Snyder, M.P., Chang, H.Y., and Greenleaf, W.J. (2015). Single-cell chromatin accessibility reveals principles of regulatory variation. Nature 523, 486–490.

Buenrostro, J.D., Corces, R., Wu, B., Schep, A.N., Lareau, C., Majeti, R., Chang, H., and Greenleaf, W. (2017). Single-cell epigenomics maps the continuous regulatory landscape of human hematopoietic differentiation. bioRxiv 109843.

Buenrostro, J.D., Corces, M.R., Lareau, C.A., Wu, B., Schep, A.N., Aryee, M.J., Majeti, R., Chang, H.Y., and Greenleaf, W.J. (2018). Integrated Single-Cell Analysis Maps the Continuous Regulatory Landscape of Human Hematopoietic Differentiation. Cell 173, 1535-1548.e16.

Butler, A., Hoffman, P., Smibert, P., Papalexi, E., and Satija, R. (2018). Integrating single-cell transcriptomic data across different conditions, technologies, and species. Nat. Biotechnol. 36, 411–420.

Chen, B., Gilbert, L.A., Cimini, B.A., Schnitzbauer, J., Zhang, W., Li, G.-W., Park, J., Blackburn, E.H., Weissman, J.S., Qi, L.S., et al. (2013). Dynamic imaging of genomic loci in living human cells by an optimized CRISPR/Cas system. Cell 155, 1479–1491.

Chen, X., Litzenburger, U., Wei, Y., Schep, A.N., LaGory, E.L., Choudhry, H., Giaccia, A.J., Greenleaf, W.J., and Chang, H. (2018a). Joint single-cell DNA accessibility and protein epitope profiling reveals environmental regulation of epigenomic heterogeneity. BioRxiv 310359.

Chen, X., Natarajan, K.N., and Teichmann, S.A. (2018b). A rapid and robust method for single cell chromatin accessibility profiling. bioRxiv 309831.

Cho, S.W., Xu, J., Sun, R., Mumbach, M.R., Carter, A.C., Chen, Y.G., Yost, K.E., Kim, J., He, J., Nevins, S.A., et al. (2018). Promoter of lncRNA Gene PVT1 Is a Tumor-Suppressor DNA Boundary Element. Cell 173, 1398-1412.e22.

Corces, M.R., Buenrostro, J.D., Wu, B., Greenside, P.G., Chan, S.M., Koenig, J.L., Snyder, M.P., Pritchard, J.K., Kundaje, A., Greenleaf, W.J., et al. (2016). Lineage-specific and single-cell chromatin accessibility charts human hematopoiesis and leukemia evolution. Nat. Genet.

Corces, M.R., Trevino, A.E., Hamilton, E.G., Greenside, P.G., Sinnott-Armstrong, N.A., Vesuna, S., Satpathy, A.T., Rubin, A.J., Montine, K.S., Wu, B., et al. (2017). An improved ATAC-seq protocol reduces background and enables interrogation of frozen tissues. Nat. Methods.

Costanzo, M., Baryshnikova, A., Bellay, J., Kim, Y., Spear, E.D., Sevier, C.S., Ding, H., Koh, J.L.Y., Toufighi, K., Mostafavi, S., et al. (2010). The Genetic Landscape of a Cell. Science 327, 425–431.

Cusanovich, D.A., Daza, R., Adey, A., Pliner, H.A., Christiansen, L., Gunderson, K.L., Steemers, F.J., Trapnell, C., and Shendure, J. (2015). Multiplex single-cell profiling of chromatin accessibility by combinatorial cellular indexing.

Cusanovich, D.A., Reddington, J.P., Garfield, D.A., Daza, R.M., Aghamirzaie, D., Marco-Ferreres, R., Pliner, H.A., Christiansen, L., Qiu, X., Steemers, F.J., et al. (2018). The cis-regulatory dynamics of embryonic development at single-cell resolution. Nature 555, 538–542.

Datlinger, P., Rendeiro, A.F., Schmidl, C., Krausgruber, T., Traxler, P., Klughammer, J., Schuster, L.C., Kuchler, A., Alpar, D., and Bock, C. (2017). Pooled CRISPR screening with single-cell transcriptome readout. Nat. Methods 14, 297–301.

Dixit, A., Parnas, O., Li, B., Chen, J., Fulco, C.P., Jerby-Arnon, L., Marjanovic, N.D., Dionne, D., Burks, T., Raychowdhury, R., et al. (2016). Perturb-Seq: Dissecting Molecular Circuits with Scalable Single-Cell RNA Profiling of Pooled Genetic Screens. Cell 167, 1853-1866.e17.

Eckert, R.L., Adhikary, G., Young, C.A., Jans, R., Crish, J.F., Xu, W., and Rorke, E.A. (2013). AP1 transcription factors in epidermal differentiation and skin cancer. J. Skin Cancer 2013, 537028.

Engreitz, J.M., Haines, J.E., Perez, E.M., Munson, G., Chen, J., Kane, M., McDonel, P.E., Guttman, M., and Lander, E.S. (2016). Local regulation of gene expression by lncRNA promoters, transcription and splicing. Nature 539, 452–455.

Farh, K.K.-H., Marson, A., Zhu, J., Kleinewietfeld, M., Housley, W.J., Beik, S., Shoresh, N., Whitton, H., Ryan, R.J.H., Shishkin, A.A., et al. (2015). Genetic and epigenetic fine mapping of causal autoimmune disease variants. Nature 518, 337–343.

Feldman, D., Singh, A., Garrity, A.J., and Blainey, P.C. (2018). Lentiviral co-packaging mitigates the effects of intermolecular recombination and multiple integrations in pooled genetic screens. bioRxiv 262121.

Flynn, R.A., Do, B.T., Rubin, A.J., Calo, E., Lee, B., Kuchelmeister, H., Rale, M., Chu, C., Kool, E.T., Wysocka, J., et al. (2016). 7SK-BAF axis controls pervasive transcription at enhancers. Nat. Struct. Mol. Biol. 23, 231–238.

Guttormsen, J., Koster, M.I., Stevens, J.R., Roop, D.R., Williams, T., and Winger., Q.A. (2008). Disruption of epidermal specific gene expression and delayed skin development in AP-2? mutant mice. Dev. Biol. 317, 187–195.

Heath, J.R., Ribas, A., and Mischel, P.S. (2016). Single-cell analysis tools for drug discovery and development. Nat. Rev. Drug Discov. 15, 204–216.

Hill, A.J., McFaline-Figueroa, J.L., Starita, L.M., Gasperini, M.J., Matreyek, K.A., Packer, J., Jackson, D., Shendure, J., and Trapnell, C. (2018). On the design of CRISPR-based single-cell molecular screens. Nat. Methods 15, 271–274.

Jaitin, D.A., Weiner, A., Yofe, I., Lara-Astiaso, D., Keren-Shaul, H., David, E., Salame, T.M., Tanay, A., van Oudenaarden, A., and Amit, I. (2016). Dissecting Immune Circuits by Linking CRISPR-Pooled Screens with Single-Cell RNA-Seq. Cell 167, 1883-1896.e15.

Kaikkonen, M.U., Spann, N.J., Heinz, S., Romanoski, C.E., Allison, K.A., Stender, J.D., Chun, H.B., Tough, D.F., Prinjha, R.K., Benner, C., et al. (2013). Remodeling of the enhancer landscape during macrophage activation is coupled to enhancer transcription. Mol. Cell 51, 310–325.

Kishibe, M., Bando, Y., Terayama, R., Namikawa, K., Takahashi, H., Hashimoto, Y., Ishida-Yamamoto, A., Jiang, Y.-P., Mitrovic, B., Perez, D., et al. (2007). Kallikrein 8 Is Involved in Skin Desquamation in Cooperation with Other Kallikreins. J. Biol. Chem. 282, 5834–5841.

Klein, A.M., Mazutis, L., Akartuna, I., Tallapragada, N., Veres, A., Li, V., Peshkin, L., Weitz, D.A., and Kirschner, M.W. (2015). Droplet barcoding for single-cell transcriptomics applied to embryonic stem cells. Cell 161, 1187–1201.

Kreslavsky, T., Vilagos, B., Tagoh, H., Poliakova, D.K., Schwickert, T.A., Wöhner, M., Jaritz, M., Weiss, S., Taneja, R., Rossner, M.J., et al. (2017). Essential role for the transcription factor Bhlhe41 in regulating the development, self-renewal and BCR repertoire of B-1a cells. Nat. Immunol. 18, 442–455.

Kretz, M., Siprashvili, Z., Chu, C., Webster, D.E., Zehnder, A., Qu, K., Lee, C.S., Flockhart, R.J., Groff, A.F., Chow, J., et al. (2013). Control of somatic tissue differentiation by the long non-coding RNA TINCR. Nature 493, 231–235.

Langmead, B., and Salzberg, S.L. (2012). Fast gapped-read alignment with Bowtie 2. Nat. Methods 9, 357–359.

Lara-Astiaso, D., Weiner, A., Lorenzo-Vivas, E., Zaretsky, I., Jaitin, D.A., David, E., Keren-Shaul, H., Mildner, A., Winter, D., Jung, S., et al. (2014). Immunogenetics. Chromatin state dynamics during blood formation. Science 345, 943–949.

Lee, N., Moss, W.N., Yario, T.A., and Steitz, J.A. (2015). EBV noncoding RNA binds nascent RNA to drive host PAX5 to viral DNA. Cell 160, 607–618.

Liu, L., Liu, C., Wu, L., Quintero, A., Yuan, Y., Wang, M., Cheng, M., Xu, L., Dong, G., Li, R., et al. (2018). Deconvolution of single-cell multi-omics layers reveals regulatory heterogeneity. bioRxiv 316208.

Liu, P., Keller, J.R., Ortiz, M., Tessarollo, L., Rachel, R.A., Nakamura, T., Jenkins, N.A., and Copeland, N.G. (2003). Bcl11a is essential for normal lymphoid development. Nat. Immunol. 4, 525–532.

Lopez, R.G., Garcia-Silva, S., Moore, S.J., Bereshchenko, O., Martinez-Cruz, A.B., Ermakova, O., Kurz, E., Paramio, J.M., and Nerlov, C. (2009). C/EBPalpha and beta couple interfollicular keratinocyte proliferation arrest to commitment and terminal differentiation. Nat. Cell Biol. 11, 1181–1190.

Lopez-Pajares, V., Qu, K., Zhang, J., Webster, D.E., Barajas, B.C., Siprashvili, Z., Zarnegar, B.J., Boxer, L.D., Rios, E.J., Tao, S., et al. (2015). A LncRNA-MAF:MAFB transcription factor network regulates epidermal differentiation. Dev. Cell 32, 693–706.

Lunning, M.A., and Green, M.R. (2015). Mutation of chromatin modifiers; an emerging hallmark of germinal center B-cell lymphomas. Blood Cancer J. 5, e361.

Ma, S., Pathak, S., Trinh, L., and Lu, R. (2008). Interferon regulatory factors 4 and 8 induce the expression of Ikaros and Aiolos to down-regulate pre–B-cell receptor and promote cell-cycle withdrawal in pre–B-cell development. Blood 111, 1396–1403.

McKercher, S.R., Torbett, B.E., Anderson, K.L., Henkel, G.W., Vestal, D.J., Baribault, H., Klemsz, M., Feeney, A.J., Wu, G.E., Paige, C.J., et al. (1996). Targeted disruption of the PU.1 gene results in multiple hematopoietic abnormalities. EMBO J. 15, 5647–5658.

Medzhitov, R., and Horng, T. (2009). Transcriptional control of the inflammatory response. Nat. Rev. Immunol. 9, 692–703.

Mezger, A., Klemm, S., Mann, I., Brower, K., Mir, A., Bostick, M., Farmer, A., Fordyce, P., Linnarsson, S., and Greenleaf, W. (2018). High-throughput chromatin accessibility profiling at single-cell resolution. bioRxiv 310284.

Mumbach, M.R., Satpathy, A.T., Boyle, E.A., Dai, C., Gowen, B.G., Cho, S.W., Nguyen, M.L., Rubin, A.J., Granja, J.M., Kazane, K.R., et al. (2017). Enhancer connectome in primary human cells identifies target genes of disease-associated DNA elements. Nat. Genet.

Najm, F.J., Strand, C., Donovan, K.F., Hegde, M., Sanson, K.R., Vaimberg, E.W., Sullender, M.E., Hartenian, E., Kalani, Z., Fusi, N., et al. (2018). Orthologous CRISPR–Cas9 enzymes for combinatorial genetic screens. Nat. Biotechnol. 36, 179–189.

Nutt, S.L., and Kee, B.L. (2007). The Transcriptional Regulation of B Cell Lineage Commitment. Immunity 26, 715–725.

Pang, S.H.M., Minnich, M., Gangatirkar, P., Zheng, Z., Ebert, A., Song, G., Dickins, R.A., Corcoran, L.M., Mullighan, C.G., Busslinger, M., et al. (2016). PU.1 cooperates with IRF4 and IRF8 to suppress pre-B-cell leukemia. Leukemia 30, 1375–1387.

Qiu, X., Mao, Q., Tang, Y., Wang, L., Chawla, R., Pliner, H.A., and Trapnell, C. (2017). Reversed graph embedding resolves complex single-cell trajectories. Nat. Methods 14, 979–982.

Rada-Iglesias, A., Bajpai, R., Swigut, T., Brugmann, S.A., Flynn, R.A., and Wysocka, J. (2011). A unique chromatin signature uncovers early developmental enhancers in humans. Nature 470, 279–283.

van Riel, B., and Rosenbauer, F. (2014). Epigenetic control of hematopoiesis: the PU.1 chromatin connection. Biol. Chem. 395, 1265–1274.

Roadmap Epigenomics Consortium, Kundaje, A., Meuleman, W., Ernst, J., Bilenky, M., Yen, A., Heravi-Moussavi, A., Kheradpour, P., Zhang, Z., Wang, J., et al. (2015). Integrative analysis of 111 reference human epigenomes. Nature 518, 317–330.

Rubin, A.J., Barajas, B.C., Furlan-Magaril, M., Lopez-Pajares, V., Mumbach, M.R., Howard, I., Kim, D.S., Boxer, L.D., Cairns, J., Spivakov, M., et al. (2017). Lineage-specific dynamic and pre-established enhancer-promoter contacts cooperate in terminal differentiation. Nat. Genet. 49, 1522–1528.

Samanta, M., Iwakiri, D., Kanda, T., Imaizumi, T., and Takada, K. (2006). EB virus-encoded RNAs are recognized by RIG-I and activate signaling to induce type I IFN. EMBO J. 25, 4207–4214.

Sanjana, N.E., Shalem, O., and Zhang, F. (2014). Improved vectors and genome-wide libraries for CRISPR screening. Nat. Methods 11, 783–784.

Satpathy, A.T., Saligrama, N., Buenrostro, J.D., Wei, Y., Wu, B., Rubin, A.J., Granja, J.M., Lareau, C.A., Li, R., Qi, Y., et al. (2018). Transcript-indexed ATAC-seq for precision immune profiling. Nat. Med. 24, 580–590.

Schep, A.N., Buenrostro, J.D., Denny, S.K., Schwartz, K., Sherlock, G., and Greenleaf, W.J. (2015). Structured nucleosome fingerprints enable high-resolution mapping of chromatin architecture within regulatory regions. Genome Res. 25, 1757–1770.

Schep, A.N., Wu, B., Buenrostro, J.D., and Greenleaf, W.J. (2017). chromVAR: inferring transcription-factor-associated accessibility from single-cell epigenomic data. Nat. Methods 14, 975–978.

Scott, E.W., Simon, M.C., Anastasi, J., and Singh, H. (1994). Requirement of transcription factor PU.1 in the development of multiple hematopoietic lineages. Science 265, 1573–1577.

Scott, E.W., Fisher, R.C., Olson, M.C., Kehrli, E.W., Simon, M.C., and Singh, H. (1997). PU.1 functions in a cell-autonomous manner to control the differentiation of multipotential lymphoid-myeloid progenitors. Immunity 6, 437–447.

Sen, G.L., Boxer, L.D., Webster, D.E., Bussat, R.T., Qu, K., Zarnegar, B.J., Johnston, D., Siprashvili, Z., and Khavari, P.A. (2012). ZNF750 is a p63 target gene that induces KLF4 to drive terminal epidermal differentiation. Dev. Cell 22, 669–677.

Stehling-Sun, S., Dade, J., Nutt, S.L., DeKoter, R.P., and Camargo, F.D. (2009). Regulation of lymphoid versus myeloid fate “choice” by the transcription factor Mef2c. Nat. Immunol. 10, 289–296.

Su, G.H., Ip, H.S., Cobb, B.S., Lu, M.M., Chen, H.M., and Simon, M.C. (1996). The Ets protein Spi-B is expressed exclusively in B cells and T cells during development. J. Exp. Med. 184, 203–214.

Tamura, T., Yanai, H., Savitsky, D., and Taniguchi, T. (2008). The IRF family transcription factors in immunity and oncogenesis. Annu. Rev. Immunol. 26, 535–584.

Vierstra, J., Rynes, E., Sandstrom, R., Zhang, M., Canfield, T., Hansen, R.S., Stehling-Sun, S., Sabo, P.J., Byron, R., Humbert, R., et al. (2014). Mouse regulatory DNA landscapes reveal global principles of cis-regulatory evolution. Science 346, 1007–1012.

Xie, S., Cooley, A., Armendariz, D., Zhou, P., and Hon, G.C. (2018). Frequent sgRNA-barcode recombination in single-cell perturbation assays. PLoS ONE 13, e0198635.

Yang, X., Lu, H., Yan, B., Romano, R.-A., Bian, Y., Friedman, J., Duggal, P., Allen, C., Chuang, R., Ehsanian, R., et al. (2011). ?Np63 versatilely regulates a broad NF-?B gene program and promotes squamous epithelial proliferation, migration and inflammation. Cancer Res. 71, 3688–3700.

Zhang, Y., Liu, T., Meyer, C.A., Eeckhoute, J., Johnson, D.S., Bernstein, B.E., Nusbaum, C., Myers, R.M., Brown, M., Li, W., et al. (2008). Model-based analysis of ChIP-Seq (MACS). Genome Biol. 9, R137.

Zhao, B., Barrera, L.A., Ersing, I., Willox, B., Schmidt, S.C.S., Greenfeld, H., Zhou, H., Mollo, S.B., Shi, T.T., Takasaki, K., et al. (2014). The NF-?B Genomic Landscape in Lymphoblastoid B-cells. Cell Rep. 8, 1595–1606.

